# Mapping human pressures across the planet uncovers anthropogenic threat complexes

**DOI:** 10.1101/432880

**Authors:** D.E. Bowler, A.D. Bjorkman, M. Dornelas, I.H. Myers-Smith, L. M. Navarro, A. Niamir, S.R. Supp, C. Waldock, M. Vellend, S. A. Blowes, K. Böhning-Gaese, H. Bruelheide, R. Elahi, L.H. Antão, J. Hines, F. Isbell, H.P. Jones, A.E. Magurran, J. S. Cabral, M. Winter, A.E. Bates

## Abstract

1. Climate change and other anthropogenic drivers of biodiversity change are unequally distributed across the world. Overlap in the distributions of different drivers have important implications for biodiversity change attribution and the potential for interactive effects. However, the spatial relationships among different drivers, and whether they differ between the terrestrial and marine realm has yet to be examined.
2. We compiled global gridded datasets on climate change, land-use, resource exploitation, pollution, alien species potential, and human population density. We used multivariate statistics to examine the spatial relationships amongst the drivers and to characterize the typical combinations of drivers experienced by different regions of the world.
3. We found stronger positive correlations among drivers in the terrestrial than in the marine realm, leading to areas with high intensities of multiple drivers on land. Climate change tended to be negatively correlated with other drivers in the terrestrial realm (e.g., in the tundra and boreal forest with high climate change but low human use and pollution), whereas the opposite was true in the marine realm (e.g., in the Indo-Pacific with high climate change and high fishing).
4. We show that different regions of the world can be defined by anthropogenic threat complexes (ATCs), distinguished by different sets of drivers with varying intensities. The ATCs can be used to test hypotheses about patterns of biodiversity change, especially in response to the joint effects of multiple drivers. More generally, our global analysis highlights the broad conservation priorities needed to mitigate the impacts of anthropogenic change, with different priorities emerging on land and in the ocean, and in different parts of the world.

## Introduction

Human activities are reshaping biological communities and impacting ecosystem functioning across the Earth (Barnosky *et al*. 2011; Dornelas *et al*. 2014; Isbell *et al*. 2017). Meeting the global challenge of the conservation and sustainable use of nature requires not only quantifying biodiversity change, but also identifying the underlying causes of change (Tittensor *et al*. 2014; Isbell *et al*. 2017). Climate change, habitat change, exploitation, pollution and invasive alien species have been recognized as the most important and widespread direct causes of biodiversity change (Pereira, Navarro & Martins 2012; IPCC 2013; IPBES 2019). These five main drivers have been linked with changes in multiple dimensions of biodiversity, including genetic diversity, species’ population sizes and the functional diversity of communities (Pereira, Navarro & Martins 2012). The impacts of these drivers on a biological community in a given region critically hinge on the extent of exposure to each driver, which refers to its local magnitude or change (such as the strength of climate change or intensity of land-use). An important, but so far underexplored, step towards understanding the global patterns of biodiversity change is characterizing the exposure patterns of biological communities to different types of environmental change.

Global maps of pressures such as the terrestrial human footprint (Sanderson *et al*. 2002; Venter *et al*. 2016), marine pressures (Halpern *et al*. 2008; Halpern *et al*. 2015a) and river threats (Vorosmarty *et al*. 2010) characterize the geographic hotspots of anthropogenic threats to biodiversity. These maps have estimated that at least 75% of terrestrial land has been exposed to some sort of land-use change (Venter *et al*. 2016), while the marine realm has been exposed to multiple pressures (Halpern *et al*. 2015a). The intensity of the terrestrial human footprint has been linked with spatial variation in ecological processes; for instance, reduced animal movement was found in areas with a higher human footprint across different species (Tucker *et al*. 2018). However, these global maps show the summed pressure of different drivers related to human activities and ignore any relationships among them. Areas identified as having high human pressure could be underlaid by different combinations of drivers with varying intensity, each of which may have contrasting impacts on biodiversity. Hence, while useful, these cumulative threat maps do not help understand the relative contribution of each driver to global biodiversity change, nor the potential for interactive impacts among drivers.

The relative importance of different drivers for biodiversity change and ecosystem services is a key component of both policy-oriented assessments such as IPBES framework (Diaz *et al*. 2015) and conservation targets such as CBD Aichi Biodiversity Targets (Tittensor *et al*. 2014). Hence, unpacking the spatial patterns of exposure of different drivers, and assessing the extent to which they jointly act on communities, is an important line of research. Many studies have examined the relationships between different anthropogenic drivers and biodiversity change, such as change in biological communities (richness, composition and population abundances), but they usually focus on only one or two drivers, such as climate change and land use change (Sirami *et al*. 2017). Multiple drivers may be associated with each other, either coincidentally (due to shared causes) or causally (when one driver affects the intensity of another) (Geary *et al*. 2019). Moreover, when drivers co-occur, their impacts on communities may be additive, or interact synergistically (total impact stronger when together) or antagonistically (total impact weaker when together) (Cote, Darling & Brown 2016; Sirami *et al*. 2017). Nonetheless, few studies have examined the effects of multiple drivers on biodiversity (Sirami *et al*. 2017; Mazor *et al*. 2018).

Here, we analyzed the spatial relationships among variables related to the main drivers of biodiversity change, and show how they overlap in different biogeographic regions across the entire surface of the world. The motivation for our study was primarily to advance the study of the attribution of biodiversity change, especially regarding potential drivers of long-term changes in species’ populations and communities (Dornelas *et al*. 2014; Bowler *et al*. 2017; Dornelas *et al*. 2019). Hence, our questions were focused towards the relevance of the spatial patterns of drivers for biodiversity change rather than on explaining the spatial patterns in drivers themselves. We determined the extent of driver overlap to assess the potential for different drivers to either act alone on communities, rendering their specific fingerprints easier to detect, or for drivers to act in combination, with the potential for interactive effects. For many drivers, it can be hypothesized that exposure patterns may be inter-linked due to related local or regional human activities, driven by local human population density (Ellis *et al*. 2010; Geary *et al*. 2019). In contrast, climate change is expected to be distributed differently to other variables because it is an outcome of processes at regional and global scales (IPCC 2013).

Towards these aims, we compiled global spatial gridded datasets on variables that characterize different anthropogenic drivers (Tables 1 and S1–S2). Although the specific variables differ among realms, we aligned each variable to the dominant drivers that are common across both realms (Table 1). We quantified the strengths of the spatial relationships among the intensities of the different variables related to climate change, habitat conversion and exploitation (grouped together as ‘human use’), pollution and potential for alien species immigration. Based on these relationships, we defined ‘anthropogenic threat complexes’ that typify the combinations of drivers impacting different regions of the world. Studies mapping drivers of biodiversity change have so far considered the terrestrial and marine realms separately. By employing a standardized analysis for both realms, our study also highlights similarities and differences in anthropogenic environmental changes across the world, including across realms.

**Table 1.** Anthropogenic drivers of biodiversity change and their respective variables based on available global spatial datasets (Tables S1 and S2). Variables in the same line do not necessarily represent the equivalent variable in each realm.

| Anthropogenic driver of biodiversity change | Associated variables |  |
| --- | --- | --- |
|  | Terrestrial | Marine |
| **Climate Change** | Temperature trend<br>Temperature divergence<br>Change in climate extremes<br>Velocity of climate change<br>Aridity trend | Temperature trend<br>Temperature divergence<br>Change in climate extremes<br>Velocity of climate change<br>Ocean acidification |
| **Human use**<br>**(land/sea use or change,**<br>**resource extraction,**<br>**exploitation)** | Crop cover<br>Pasture cover<br>Urban cover<br>Forest loss<br>Livestock density | Destructive demersal fishing<br>Low by-catch demersal fishing<br>High by-catch demersal fishing<br>Low by-catch pelagic fishing<br>High by-catch pelagic fishing |
| **Human population density** | Population density | Coastal population density |
| **Pollution** | Atmospheric nitrogen deposition<br>Nitrogen fertilizer application<br>Pesticide application<br>Light pollution | Ocean pollution<br>Fertilizer coastal pollution<br>Pesticide coastal pollution<br>Light pollution |
| **Alien potential** | Connectivity (transport infrastructure) | Port cargo volume |

## Methods

### Approach to data selection

By drivers, we refer to anthropogenic drivers only and not biological processes, such as dispersal and demographic rates, that more proximately lead to changes in species’ abundances. Our analysis required datasets that were (1) global; (2) spatially gridded at high-resolution and (3) publicly available. We selected datasets on variables related to the main five drivers that were included in previous studies of global environmental change in the context of biodiversity change (Table 1) (Sanderson *et al*. 2002; Halpern *et al*. 2008). We further searched for data on other relevant variables following the IUCN threats categories (Table S1) (Salafsky *et al*. 2008). As we focused on a land versus ocean comparison, we did not specifically consider freshwater threats (Vorosmarty *et al*. 2010). Variables representing drivers of biodiversity change were in most cases based on data between 1990 and 2010 (except climate change, and forest loss; see below). In total, we had datasets for 16 driver-related variables (Table 1). The terrestrial datasets came from various sources (Table S2). Most of the marine datasets came from the landmark study of Halpern et al. (Halpern *et al*. 2008). For interpretation and presentation purposes, variables were grouped into five broad categories: climate change, habitat conversion, exploitation, pollution and alien species potential. Because habitat conversion and exploitation were difficult to classify separately across terrestrial and marine ecosystems, we combined both into a single “human use” category.

### Climate change

Climate change has multiple components (IPCC 2013), hence we characterized climate change by several variables using global spatiotemporal gridded temperature data for the terrestrial (Harris *et al*. 2014) and marine realm (Rayner *et al*. 2003). To calculate temperature trends, we used data between 1950, proposed as the start of the Anthropocene (Waters *et al*. 2016), and 2010. Temperature trends were estimated by fitting simple linear regression models to annual temperature means of each grid cell and extracting the coefficient for the effect of year. Temperature divergence was inferred from the t-statistic of this linear regression, i.e., the trend divided by its standard error, representing the significance of the trend. Velocity of climate change (Loarie *et al*. 2009) was calculated as the ratio between the temporal temperature trend and the local spatial gradient in temperature. Trends of extreme temperatures were calculated by whichever was largest of the temporal trends in mean temperature of the warmest or coolest month. We further included aridity for the terrestrial realm and ocean acidification for the marine realm. The aridity trend was estimated by taking monthly and annual datasets on potential evapotranspiration and precipitation, and calculating the monthly:annual ratio (Zorner *et al*. 2008), followed by the temporal trend of the annual monthly average of this ratio. Ocean acidification, from the Halpern layers, was based on the change in aragonite saturation state between 1870 and 2000–2009 (Halpern *et al*. 2008).

### Human use

In the terrestrial realm, we collated human use variables related to different types of land conversion or use: cropland, pasture land, cattle density, urban area and forest loss. Data on crop land, pasture land and urban/built land-cover were taken from different databases, primarily based on satellite imagery – crop land (Fritz *et al*. 2015), pasture land (Ramankutty *et al*. 2008) and urban land (Friedl *et al*. 2010). We additionally included information on forest loss since deforestation itself is a recognized threat (Barlow *et al*. 2016). Forest loss, based on FAO (Food and Agriculture Association of the United Nations) wood harvest statistics, was calculated as the loss of primary forest for the same time frame as our climate change statistics, i.e., between 1950 and 2010 (Hurtt *et al*. 2011). We also included data on cattle density, which was based on sub-national livestock data that were statistically downscaled using multiple predictors (Robinson *et al*. 2014). In the marine realm, human use variables were based on different commercial fishing activities separated by gear types (e.g., dredging or cast nets), which determine their selectivity and impact on the surrounding seascape (Halpern *et al*. 2008). These fishing types were pelagic low-bycatch, pelagic high-bycatch, demersal habitat-modifying, demersal non-habitat-modifying low-bycatch, and demersal non-habitat-modifying high bycatch. These data were based on FAO and other commercial catch data sources and downscaled based on an ocean productivity model (Halpern *et al*. 2008).

### Pollution

Nitrogen from both fossil fuel combustion and agriculture is one is the biggest pollutants impacting biodiversity (De Schrijver *et al*. 2011). We included data on nitrogen pollution for the terrestrial realm in the form of atmospheric nitrogen (Dentener 2006) and fertilizer use (Potter *et al*. 2010), and for the marine realm as fertilizer use (Halpern *et al*. 2008). We also included data on pesticide use in both realms (Halpern *et al*. 2008; Vorosmarty *et al*. 2010), another important component of agricultural intensification that negatively affects biodiversity (Geiger *et al*. 2010). Country-specific estimates of fertilizer use and pesticide were downscaled to a raster grid by the data providers according to crop land-cover maps; thus, these datasets were not fully independent of the cropland data, but they still represent the best current knowledge of the spatial distribution of these variables. We also included a layer estimating the extent of ocean pollution, based on the distribution of shipping lanes (Halpern *et al*. 2008). Finally, we included night-time light pollution detected by satellite imagery (Halpern *et al*. 2015a), which was also included in previous terrestrial and marine threat maps (Halpern *et al*. 2015a; Venter *et al*. 2016).

### Alien species potential

The spread of alien species is among the greatest threats to biodiversity and ecosystem services (Blackburn, Bellard & Ricciardi 2019). Alien species are defined as species that are introduced into areas beyond their historical range, usually through human transport, accidentally or incidentally (Hulme 2009; Seebens *et al*. 2015). We used information on transport infrastructure related to human movement and trade that depict possible species transportation pathways and vectors (Hulme 2009; Davidson *et al*. 2018). Specifically, we used spatial datasets of transport connectivity (including data on road and rail networks and navigable rivers) in the terrestrial realm and cargo volume at ports in the marine realm (Table S1). While these do not represent the only invasion pathways, they are commonly accepted proxies for human-mediated propagule pressure, which is known to be among the most important determinant of alien species establishment (Hulme 2009; Seebens *et al*. 2015).

We note two limitations of our alien species data. First, our proxy was for alien species in general and not specifically for invasive and harmful alien species. Only a proportion of alien species become invasive, but this proportion differs among taxa (Jeschke & Pyšek 2018). Although the most harmful alien species have been identified for some regions (Katsanevakis *et al*. 2014), this is an on-going process at the global level (Pagad *et al*. 2018). Hence, we refer to this variable as alien species potential rather than specifically as invasive alien species. Second, we only used a proxy for alien species propagule pressure and not direct measures of their distribution. Our analysis required high-resolution global gridded maps, but these are not available for alien species distributions for neither realm. However, information on alien species distribution was available at a regional, sub-national and national levels for some taxonomic groups, including birds (Dyer *et al*. 2017) and plants (van Kleunen *et al*. 2019). To assess the validity of our connectivity proxy, we used these datasets, representing taxa with low and high mobility, to test the correlation between alien species richness and mean connectivity at the spatial scale of the distribution data. We found a significant rank correlation for both datasets (birds, ρ=0.42; plants, ρ=0.46, see Fig. S1 for more details), suggesting our proxy represents a reasonable approximation of alien species potential.

### Human population

We also included “human population density” as a separate driver (CIESIN 2017) accounting for the effects of human activities not falling into the other categories (Salafsky *et al*. 2008). Including human population density also allowed quantitative tests of the relationship between human population density and the other drivers.

### Justification for layer exclusion

We did not use data for some variables that were previously included in the terrestrial human footprint or the Halpern layers. The human footprint includes data on roads, railways and navigable waterways (Venter *et al*. 2016). Although we did not separately include these data, these data has been taken into account in the terrestrial connectivity variable (for invasions). In the marine realm, we excluded a shipping lane variable since the ocean pollution variable was already based on the distribution of shipping lanes (Halpern *et al*. 2008). Additional available marine layers that we did not use were: UV radiation, oils rigs (based on night lights, already included), inorganic pollution (highly correlated with other land-based coastal pollutants that were already included) and artisanal fishing (data poor and mostly modelled) (Halpern *et al*. 2008).

### Geographic region data

To understand how the driver distributions compared with classic biome distributions that are already used in biodiversity research, we also extracted geographic region data on biomes. Data on the spatial distribution of terrestrial biomes (including deserts, temperate broadleaf forest and boreal forest among other) were taken from WWF (Olson *et al*. 2001). Marine regions were defined by combining coastal/shelf region polygon data – MEOW (Spalding *et al*. 2007) and ocean polygon data (naturalearthdata.com). We did not use marine ecosystem data as used by others (Halpern *et al*. 2015a) because the ecosystems spatially overlapped in our 2-D global raster grid, when, in reality, these different ecosystems occur at different depths in the water column.

### Data processing

We harmonized each dataset to a standard global grid. The resolutions of the original datasets were approximately at a 100 km square grid (or 1°) or finer resolution; hence, we aggregated all datasets to a standard grid of 100 km square grid cells by taking the mean value of the grid cells. Atmospheric nitrogen deposition was only available at a coarser resolution (see Table S1); however, we disaggregated this also to 100 km. Datasets were bound between latitudes of −58 and 78 (due to poorer data at extreme latitudes) and re-projected onto a common equal-area map projection (Eckert IV; ESPG = 54012). Missing values in some of the human activity datasets (primarily in pesticide use) occurred in remote regions (e.g., very high latitudes and deserts) with likely absent or low variable values (since they were in areas with no crop cover) and were imputed as zero. Greenland was excluded due to missing data in several of the datasets.

Because each dataset comprised data in different units (e.g., temperature data in °C and fertilizer data in kg/ha), it was not possible to directly compare their absolute values. Instead, we ranked scaled (between 0 and 1) values of each dataset for ease of interpretation (Fig. S2 shows the distributions of the original values of each variable and Fig. S3 shows global maps of the ranked and scaled data). This processing also reduced the large skew in the absolute values of many datasets. For most datasets, larger values reflected a greater magnitude, and thus higher potential exposure of biodiversity to that variable. Transformations were needed in only one case – we inverted terrestrial accessibility (i.e., *values*^−1^). For climate change metrics, we only used positive values to focus on the drivers of warming and drying. By using ranks, we avoided making any complex assumptions about the relationships between the absolute values of each driver variable and its impact on organisms. We rather assumed that all variables were similarly important and that higher variable values would have a stronger impact on biodiversity.

### Data analysis

To examine the relationships among the intensities of the 16 different variables, we calculated Spearman’s rank correlation coefficients (ρ) for each pairwise combination of variables across all grid cells in each realm. We chose this statistic because it is equivalent to the commonly used Pearson’s correlation on ranked data, which was consistent with our aforementioned data processing. We used Dutilleul’s modified t-test to account for spatial autocorrelation before testing the significance of the correlations (Dutilleul, Pelletier & Alpargu 2008). We also used Moran’s I and correlograms to assess the extent of spatial autocorrelation within each variable. For the marine realm, correlations were also examined separately for grid cells whose centroid overlapped with oceanic or coastal regions. To assess the importance of the drivers in different parts of the world, we compared the average intensity of each of the five main drivers (climate change, human use, human population, pollution and alien species potential) for different geographic regions. We first calculated the mean driver value for each grid cell, i.e., averaging across associated driver-related variables (Table 1). We then plotted the distribution of these mean values across all grid cells within each terrestrial biome and marine region.

We used k-medoid clustering, with the partitioning around the medoids algorithm with Euclidean distances (Maechler *et al*. 2018), for clustering grid cells according to their extent of exposure of all the variables. To focus the clustering on the main axes of variation, we first used PCA to reduce the number of variables in driver groups with multiple variables, i.e., climate change, human use and pollution. We performed the clustering with two or three PCA components (whichever captured at least 75% of the variation) for each of these drivers along with the variables for human population density and potential alien species. We selected the number of clusters by comparing the changes in cluster silhouette width with increasing cluster number; however, we limited the cluster number to <10. To slightly smooth the maps, we used a moving window to assign each cell the mode of its 3 x 3 cell neighborhood. Although, driver combinations vary in a continuous manner, we chose a clustering method that produces discrete grouping to provide the simplest description of the main groupings in the data. Finally, to repack the datasets into cumulative driver maps, we summed the number of variables for which each grid cell was in the upper 10% of values (based on values greater than zero). Analyses were run in R v. 3.4.1 (R Core Team 2018), using the packages raster (Hijmans 2017), SpatialPack (Vallejos, Osorio & Bevilacqua 2018) and cluster (Maechler *et al*. 2018).

### Sensitivity analyses

To examine the effect of the grain size, we repeated the data processing steps except harmonized the datasets to a global grid of 500 km resolution and repeated the correlation analysis (similar results were obtained – see Fig. S4). To check the effects of ranking the data values, we repeated the data processing steps by logging the values (to the base 10) rather than ranking them, after bounding extreme values to the upper and lower 2.5% quantiles. This alternative data transformation does not affect the correlation coefficients because Spearman’s correlations only uses the ranks of the data. We repeated our remaining analysis with this alternative transformation, calculating the average variable intensities for different terrestrial and marine regions, and the clustering analysis (generally similar results were obtained – see Fig. S5 and S6). Since the distributions are still skewed after logging instead of ranking, the patterns are strongly affected by extreme values with this approach, especially in the marine realm.

## Results

We found that drivers of biodiversity change were more spatially coupled in the terrestrial than in the marine realm (Fig. 1, Fig. S7). On land, 40% of the possible pairwise relationships between variables (excluding climate change-related variables) showed positive correlation strengths of at least 0.7. Thus, terrestrial areas with high intensities of one variable also tended to have high intensities of other variables. Moreover, correlations were found between different types of drivers; for instance, high crop land-cover was associated with high pollution, high alien potential and high human population density. Conversely, in the marine realm, we found fewer correlations – only 15% of the possible pairwise relationships (excluding climate change-related variables) showed a strong positive correlation (> 0.7) – and these relationships were mostly within, rather than between, different driver types; for instance, among different types of human use (e.g., different types of demersal fishing; Fig. 1). Across all variables, oceanic regions showed fewer correlations compared to coastal regions (Fig. S8). Spatial autocorrelation was present in all variables and tended to reach greater distances in the marine human-uses and climate-change variables (Figs S9 and S10), and shorter distances in the coastal-based marine variables. However, the correlations among drivers were statistically significant (all P<0.05 for links shown in Fig. 1) after accounting for autocorrelation. In neither realm were there strong negative correlations among variables (Fig. S11 shows the full correlation matrix).

**Fig. 1.**
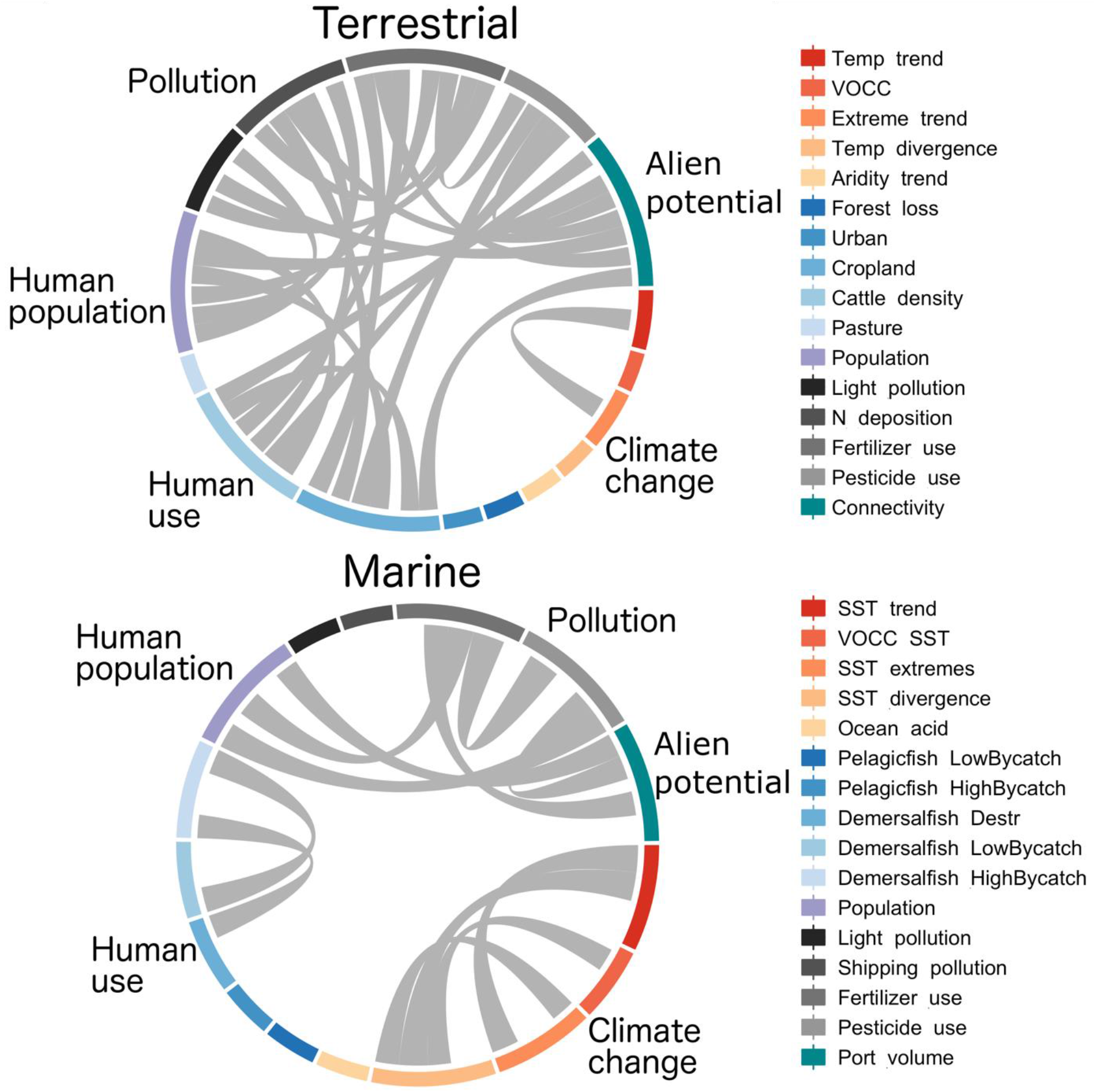
Strong and positive relationships among anthropogenic drivers of biodiversity change. Each grey link between two variables represents a significant positive correlation with strength >0.7 of the variable intensities across 100 square km grids covering each realm. We find a higher number of correlations between drivers in the terrestrial versus the marine realm.

Strong correlations between climate change and non-climatic drivers (human use, pollution and alien species) were not observed in either realm (Fig. 1, Fig. S11); however, there were still significant weak correlations, with the direction of these correlations differing systematically between realms (Fig. 2). Temperature change, and its velocity, was negatively associated with the average intensity of other variables in the terrestrial realm (Fig. 2; temperature change: ρ = −0.26, P<0.01; velocity: ρ = −0.22, P<0.01), but positively associated with the average intensity of other variables in the marine realm (Fig. 2; temperature change: ρ = 0.20, P=0.06; velocity: ρ = 0.24, P<0.05). Pollution, cattle density and human population density were especially negatively correlated with temperature change in the terrestrial realm while human use (demersal and pelagic fishing) was especially positively correlated with temperature change in the marine realm (Fig. 2).

**Fig. 2.**
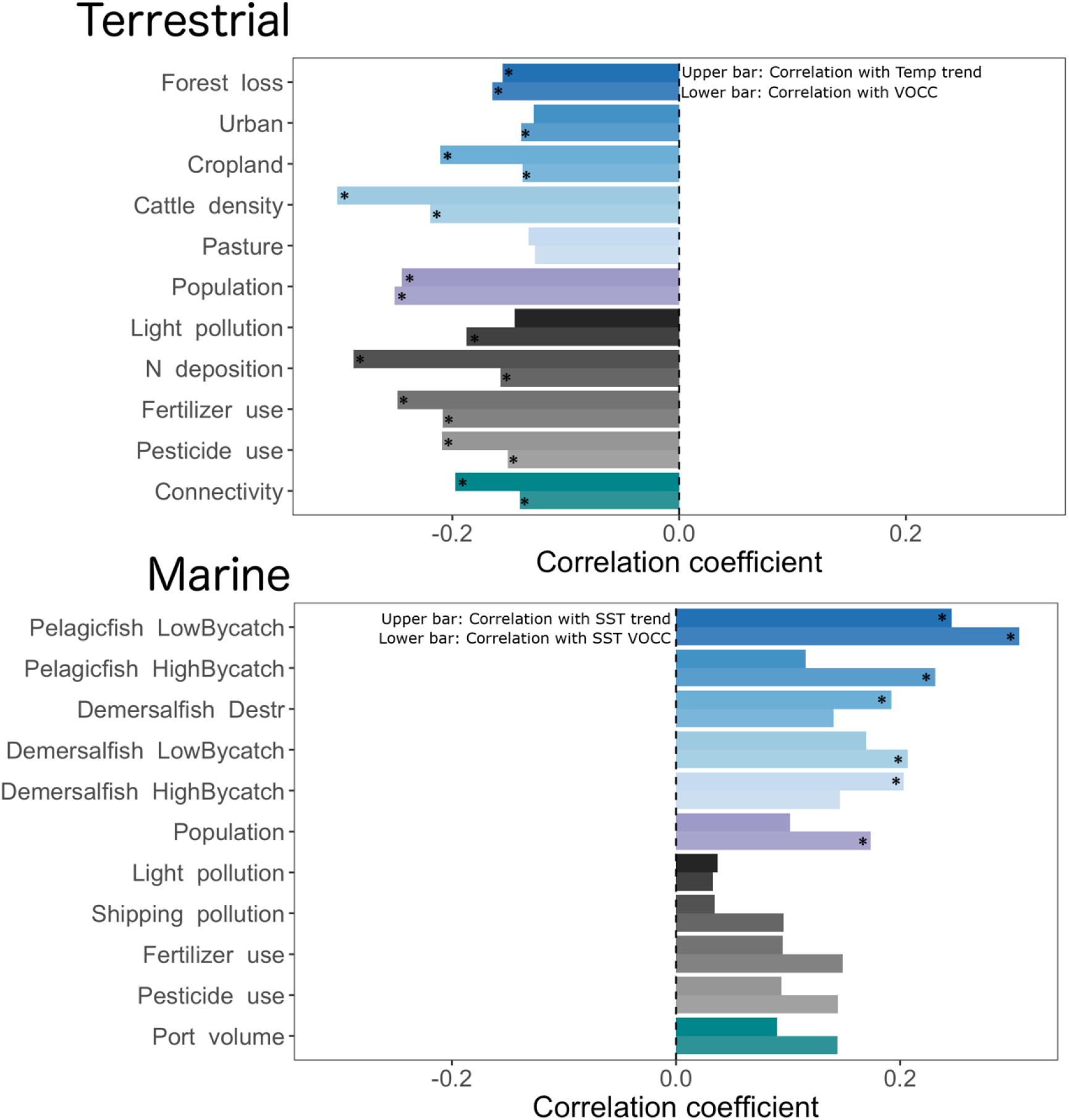
Relationships between climate change and non-climatic drivers (human use, pollution and alien species). The length of each bar represents the correlation coefficient between each variable and temperature change (upper bar, air or sea surface temperature – SST) or VOCC (lower bar, velocity of climate change). We find negative correlations in the terrestrial realm and positive correlations in the marine realm. * denotes statistical significance after accounting for spatial autocorrelation.

Consistent with the negative association between climate change and human use in the terrestrial realm, terrestrial biomes exposed to some of the strongest climate change, such as the tundra, boreal forest and desert, have experienced the lowest intensities of human use (Fig. 3, Fig. S12). By contrast, other terrestrial biomes, such as tropical dry broadleaf forest and tropical coniferous forest, have had high intensities of human use, pollution and invasions but lower intensities of climate change. Temperate broadleaf and mixed forest is distinct in the terrestrial realm by experiencing higher than average intensities of all drivers (Fig. 3). In the marine realm, central and western Indo-Pacific were most exposed to multiple drivers, including rapid climate change and multiple human uses, especially fishing (Fig. 3). The temperate Northern Atlantic, which includes the North Sea, has been also strongly exposed to multiple drivers. By contrast, the Pacific Ocean been exposed to the lowest intensities of multiple drivers.

**Fig. 3.**
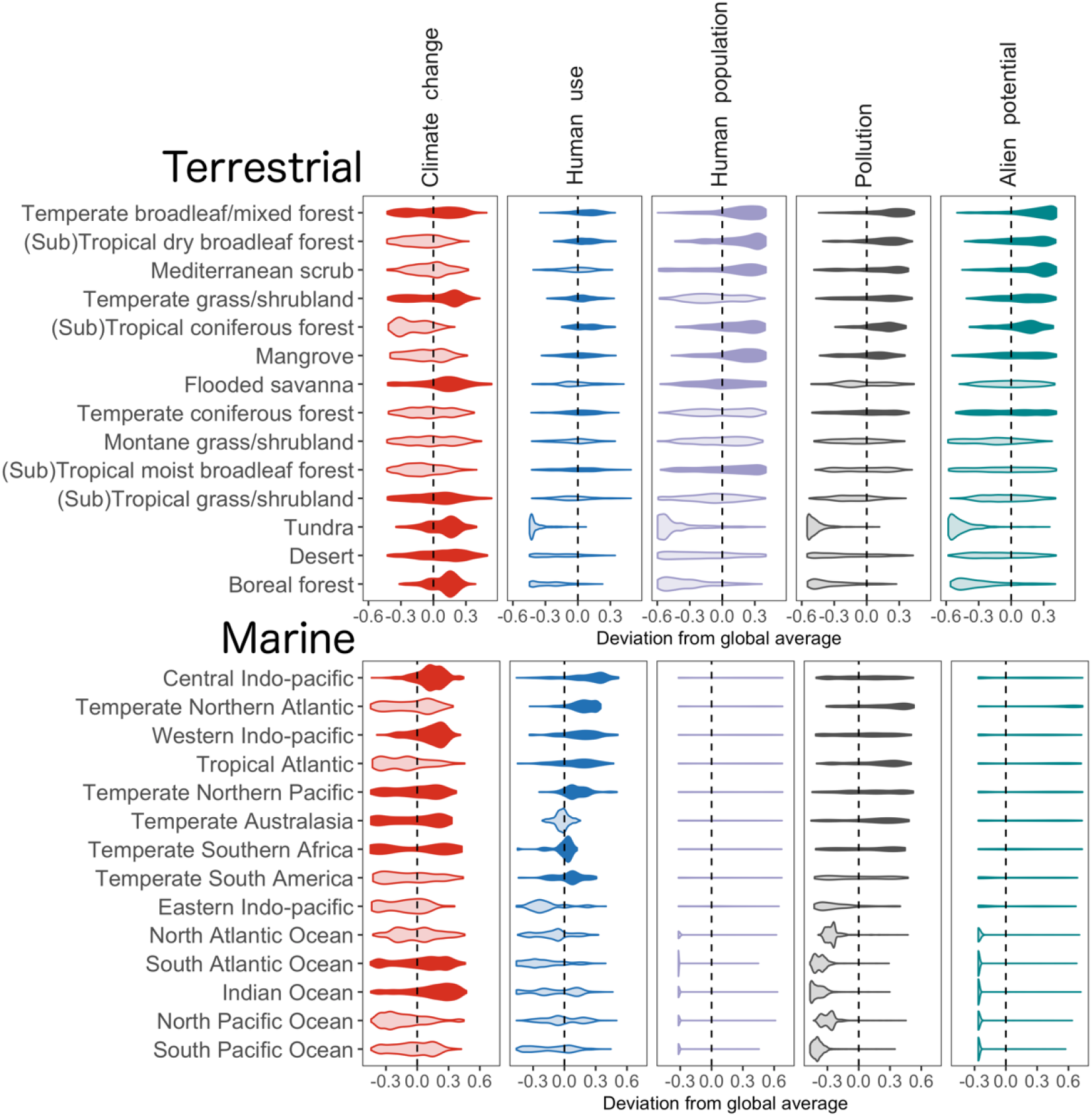
Regions of the terrestrial and marine realms are exposed to distinct combinations of drivers. The violin plots show the distribution of values for each driver in each region. Violins with a median greater than the global median of each driver (the dashed zero line; median of biome values) are colored in a darker shade. Regions are presented in declining order of total driver exposure: regions at the top are exposed to the greater intensities of drivers and those at the bottom are exposed to the least. Names of the terrestrial regions were shortened for presentation purposes. Figure S12 shows the full distributions for each individual driver variable in each region as well as gives the full names of the terrestrial regions.

The cluster analysis defined five terrestrial and six marine regions according to their similarity of exposure to the different driver variables (Fig. 4). These exposure patterns can be regarded as ‘anthropogenic threat complexes’ (ATC) that characterize the typical combinations of environmental change emerging at large scales. ATCs I and VI (dark blue-grey regions in Fig. 4) represent terrestrial and marine areas with high exposure to multiple drivers, especially non-climatic drivers such as human use, pollution and alien potential. ATCs II and VII are regions (medium blue-grey) exposed to medium intensities of many variables, including climate change. ATCs IV and IX (red regions) were similarly exposed to medium intensities of many variables, but higher intensities of climate change. ATCs III and VIII (light blue) were exposed to relatively weaker climate change than to non-climatic drivers; by contrast, ATCs V and X (orange regions) were exposed to relatively greater climate change than to non-climatic drivers. ATC XI represents areas (light orange) that were exposed to low intensities of most variables and was only found in the marine realm. The largest terrestrial ATC was V (34% of terrestrial grid cells), covering regions exposed to medium-high climate change and lower intensities of other drivers. The largest marine ATC was IX (23% of marine grid cells), including regions exposed to both medium human use, especially fishing, and medium-high climate change.

**Fig. 4.**
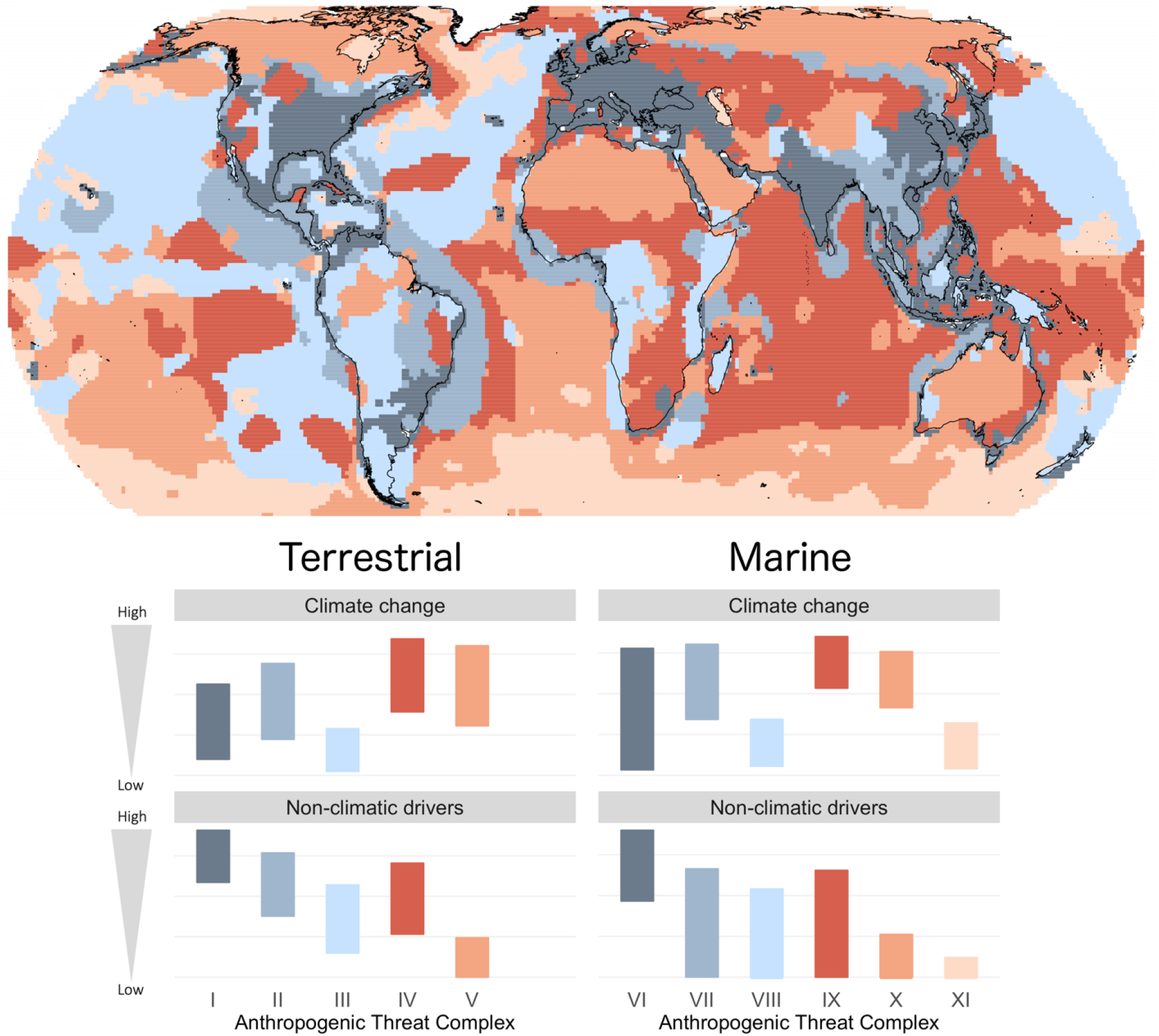
Geography of the Anthropocene. Different geographic regions of the world are exposed to different Anthropogenic Threat Complexes (labelled I to XI). In the world map, each colour in each realm denotes a region exposed to the same ATC. The bar charts below are a legend for the colours of the map that describe the magnitudes of climate change and non-climatic drivers in each ATC. Similar ATCs across realms are shaded with the same colour. The bars represent the lower and upper quartiles of the intensities of variables within each driver group (climate change or non-climatic drivers that included human use, pollution and alien species potential). Fig. S13 provides the legend showing the full suite of variable intensities in each ATC.

The global maps (Fig. 5) highlight the areas exposed to high intensities of multiple drivers and connect the ATCs to previous cumulative human impact maps produced separately for the terrestrial (Sanderson *et al*. 2002; Venter *et al*. 2016) and marine realms (Halpern *et al*. 2008; Halpern *et al*. 2015a). Terrestrial regions with the highest cumulative intensities across multiple drivers tended to be within ATC I, covering parts of Europe, Northeastern America, India as well as Northern and Eastern China (Fig. 5). In the marine realm, the highest cumulative intensities were found within ATCs VI and IX, especially in the North Sea of the Northern Atlantic, East China Sea and central Indo-Pacific. Terrestrial regions with the lowest cumulative intensities of multiple drivers were commonly within ATCs III and V, covering parts of South America, Africa, Russia and Australia. In the marine realm, the lowest cumulative intensities were within ATCs X and XI, in the Indian Ocean, the southern Atlantic Ocean and Pacific Ocean.

**Fig. 5.**
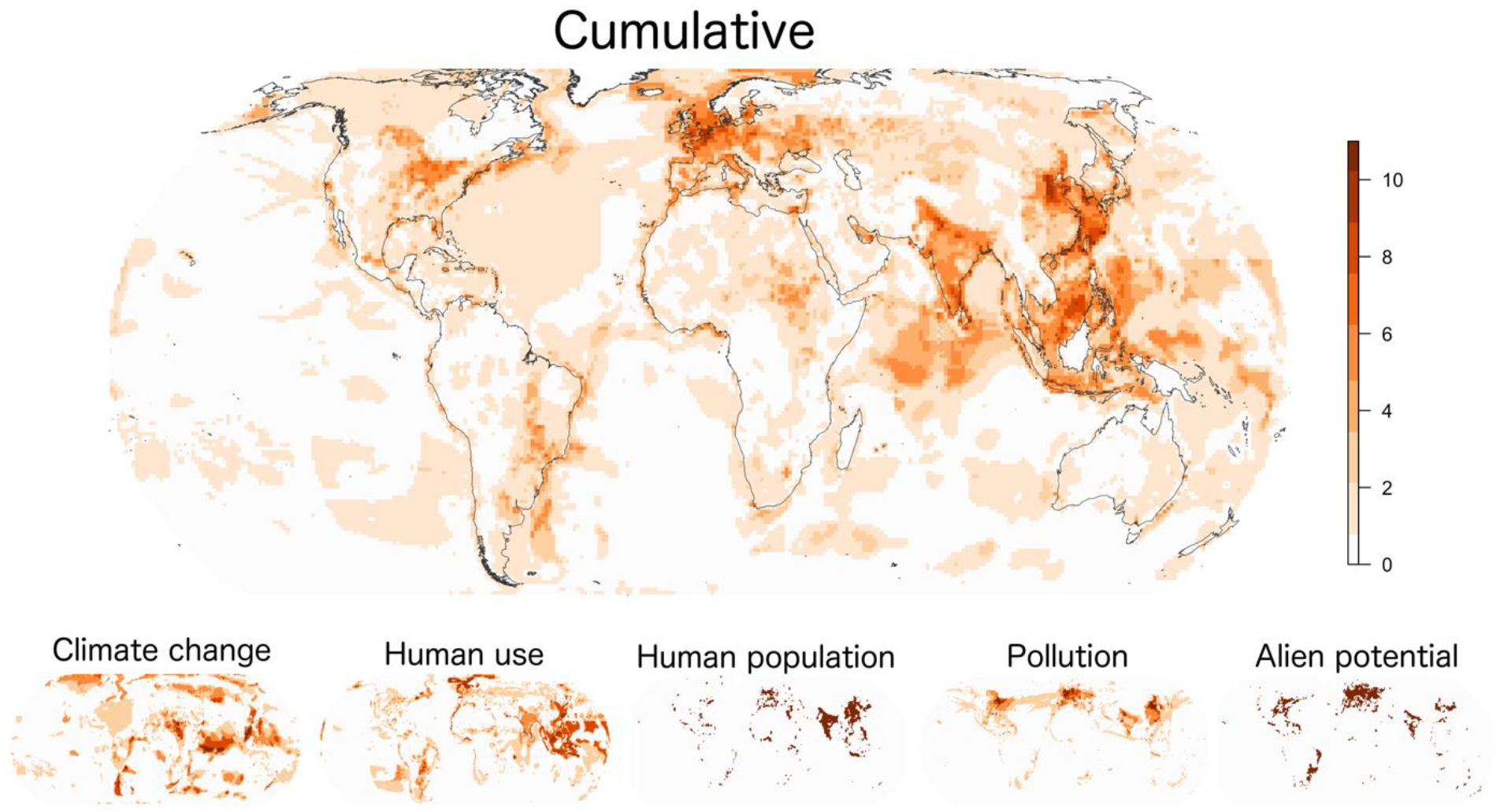
Regions of the world exposed to high intensities of multiple drivers. The main map shows the number of the 16 driver variables for which each grid cell was in the highest 10% of values within each realm. Regions in the darkest orange are exposed to high intensities of multiple variables, while those in off-white are exposed to lower intensities (i.e., within the 90% quantile of values) of all variables. The smaller plots below show the same for each of the separate drivers. Larger versions of the driver plots are presented in Fig. S14.

## Discussion

Our findings suggest that many drivers of biodiversity change are unlikely to act alone, but rather to jointly impact biological communities. In the terrestrial realm, drivers that tend to be simultaneously present in a region were associated with agricultural activities (including land conversion to cropland, use of land for cattle rearing, use of pesticides/fertilizer), human population density (urban cover and light pollution) and potential for alien species. These drivers have especially high spatial overlap in the temperate broadleaf and mixed forests of eastern North America, Europe and China. In the marine realm, drivers that tend to spatially overlap include different types of pollutants and the potential for alien species, with the coastal regions of the Indo-Pacific and Northern Atlantic (especially the North Sea) showing the highest overlap. Taken together with recent studies that highlight the lack of research on interactive effects of drivers (Sirami *et al*. 2017; Mazor *et al*. 2018), our findings emphasize the widespread relevance of further research on understanding how multiple drivers combine to impact biological communities.

Correlations among drivers were more frequent in the terrestrial realm than in the marine realm. Spatial relationships among drivers in the terrestrial realm were likely based on different activities supporting proximal human populations, leading to simultaneous pressures from land-use change and pollution (Ellis *et al*. 2010). Indeed, Venter et al. (2016) already linked spatial variation in the human footprint to land suitable for agriculture. By contrast, in the marine realm, different types of human uses were based on different fishing activities that largely occur in different areas, leading to lower spatial overlap among them. For instance, demersal fisheries, such as bottom-trawling that disturbs the seabed, mostly occurs over the continental shelf; while, pelagic fisheries, such as longline fishing for species such as tuna, can be either continental or oceanic. Coastal regions were intermediate in patterns between terrestrial and oceanic regions, suggesting that local human populations contribute to the differences between the two realms (Halpern *et al*. 2015b). Different histories of human activities in the terrestrial and marine realm likely also plays a role. Human exploitation in the terrestrial realm is more advanced, spanning millennia, while technological advances have only recently allowed greater exploitation of the world’s oceans (Knapp *et al*. 2017).

Climate change emerged from our analysis as a spatially distinct driver of biodiversity change. In both realms, climate change was only weakly associated with other drivers (human use, pollution and alien species), as expected based on the broad spatial scale at which carbon emissions affect climate. This finding suggests climate change impacts should be easier to isolate in statistical models from the impacts of other drivers, and that fingerprints of climate change impacts on species abundances, range limits and community compositions (Parmesan & Yohe 2003; Poloczanska *et al*. 2013) may be easier to detect than those of other drivers. In the terrestrial realm, climate change tended to be greater in areas with lower human use. Indeed, high-latitude terrestrial regions that are experiencing pronounced climate change (IPCC 2013; Pithan & Mauritsen 2014) have historically undergone less human settlement and agriculture. In areas where non-climatic drivers are weak, climate change has the potential to be the dominant driver of change, for instance in deserts, tundra and boreal forests of the terrestrial realm and in the south Atlantic Ocean and Indian Ocean of the marine realm.

Our classification of Anthropogenic Threat Complexes helps regard anthropogenic environmental change as a series of at least 11 large-scale natural experiments across the globe. The ATCs highlight where future research on the joint impact of multiple drivers, and their potentially interactive effects, could be most usefully directed to have the widest relevance for the study of biodiversity change. They suggest, for instance, that interactive effects between climate change and fishing should be further studied in the Indo-Pacific and North Sea, where both rapid temperature change and intense fishing have been occurring (Ramirez *et al*. 2017). In the terrestrial realm, biological communities in Europe may be of special interest to study the joint impacts of climate change, land use and pollution on different ecosystems because of the strong intensities of multiple drivers.

Attribution of observed biodiversity change to the underlying anthropogenic drivers may be most successful if focused on complexes of environmental change, rather than on each variable individually. Because of the correlations among drivers, relationships between a driver and a metric of biodiversity change may in fact be caused by a correlated driver. Although spatial heterogeneity at smaller spatial scales (e.g., neighboring sites that differ in only one driver, such as land-use change) can estimate the local effect of drivers such as habitat conversion (Newbold *et al*. 2015), correlated large-scale drivers can affect regional species pools and hence still influence local community dynamics (Harrison & Cornell 2008). Analysis of the relationship between ATCs and species’ declines may help identify the most harmful combinations of drivers. The ATCs could be resolved to finer spatial scales and also used to inform the design of biodiversity observatories that treat driver exposure as a natural experiment. Observatories could be selected along a given driver gradient, keeping all but this driver constant. Alternatively, study regions that are most suitable to study a specific driver could be selected from within geographic clusters dominated by the driver of interest, to reduce the confounding effects of other drivers in the landscape.

Large-scale information on drivers is most immediately relevant for global conservation policy (Tittensor *et al*. 2014; IPBES 2019). For biodiversity conservation at specific locations, local data on the magnitudes of different types of anthropogenic pressures are essential to decide on the most suitable management plan. However, even in this latter context, there are a number of advantages of having knowledge on the large-scale patterns of drivers. First, these large-scale patterns allow local management to be modified according to the wider anthropogenic land- or seascape context (Harrison & Cornell 2008). Whilst landscape ecology and habitat connectivity already inform conservation planning, knowledge of all drivers present in the region could influence local decision-making by enabling prediction of possible future threats and biodiversity change at local-scales, e.g., large-scale land-use change affecting regional species pools, with consequences for local community changes through dispersal (Hansen & DeFries 2007). Secondly, by characterizing regions of the world in terms of the nature of anthropogenic environmental change, the ATCs suggest how information and data might be pooled and synthesized across regions, and even across realms. Regions exposed to the same ATC, regardless of location, would benefit from exchanging knowledge about prioritization strategies and management of the multiple drivers, as well as implementing cross-border and inter-regional strategies to minimize their impact (Bonebrake *et al*. 2019).

Our findings are clearly dependent on the included layers – we selected layers based on their past inclusion in global threat maps and their linkage to IUCN threat categories. However, many of the recognized IUCN threats to biodiversity (Salafsky *et al*. 2008; Joppa *et al*. 2016) were not available as high-resolution spatial datasets at a global scale, which was necessary for our analysis. These threats include energy production and mining, hunting, and other forms of human disturbance (Salafsky *et al*. 2008). Data availability was especially limited for the marine realm and the threat from invasive alien species globally. Rather than use proxy variables of species transport and thus of propagule pressure, spatially-explicit maps of the number of invasive alien species would have improved our analysis. The Global Register of Introduced and Invasive Species (www.griis.org) is working towards better knowledge of the distribution of invasive species (Pagad *et al*. 2018). Our study only focused on spatial patterns and did not explicitly consider the consequences of changes in human activities over time. Ongoing projects such as the Copernicus project (http://www.copernicus.eu/) will greatly increase the availability of high-resolution spatiotemporal datasets on different variables in the coming years, enabling better attribution of biodiversity change to the underlying drivers.

Quantifying exposure to environmental change is the first step towards determining which species, in which places, are most impacted by human activities. However, the realized outcome of different drivers on biodiversity will ultimately depend on a combination of both the magnitude of exposure to drivers and species’ sensitivities to environmental change (Foden *et al*. 2013). While exposure characterizes the amount of environmental change (e.g., temperature change in °C), sensitivity refers to the tolerance of the biological community to that driver (e.g., abundance (or individual fitness) change per °C temperature change). We intentionally focused on exposure patterns so that our results were not species-specific and therefore potentially relevant for any taxa or ecosystem. Unlike exposure, sensitivities vary among taxa according to characteristics such as their life history, traits and niche breadth (Sunday *et al*. 2015) and therefore need to be examined separately for different taxa. Hence, despite similar exposure patterns, we can expect a diversity of biodiversity responses within each ATC due to variation in species’ sensitivities.

Our macroecological approach to mapping the drivers of biodiversity change contributes to the development of broad conservation policy targeted toward the mitigation of specific driver complexes. Development of conservation strategies that simultaneously attempt to tackle multiple drivers are likely to be more efficient in the long-term (Bonebrake *et al*. 2019). The ATC framework emphasizes the fact that multiple drivers usually affect local and regional biological communities and discourages the prevalent simplistic focus on one or two drivers. Much more research should focus on understanding the joint impacts of multiple drivers and how different drivers interact on biological communities. Our findings help direct some of these future studies by identifying which drivers most commonly overlap, and in which regions of the world. Finally, by taking a cross-realm approach, we hope to encourage information exchange across regions of the world that are exposed to similar suites of drivers, regardless of environmental realm, and the development of joined-up conservation policies across national borders and the terrestrial-marine interface.

### Data availability

Table S2 shows the sources of each dataset and links to where each dataset can be downloaded. Datasets produced during our analysis (raster layers shown in Figures 4 and 5) are available as georeferenced TIFF files in the SOM.

### Code availability

R script to harmonize the raster to a standard grid is found here: https://github.com/bowlerbear/harmonizeRasters

R script for the subsequent analysis is found here: https://github.com/bowlerbear/geographyDrivers

## Supporting information

SOM

## Acknowledgments

This paper arose from discussions at the sChange Workshop (www.idiv.de/schange), which was supported by sDiv, the synthesis centre of iDiv, the German Centre for Integrative Biodiversity Research Halle-Jena-Leipzig. We thank all other participants of this workshop for the stimulating discussion and compilation of data that directly fed into this project. We thank the Associate Editor and two anonymous reviewers for comments on the manuscript. We also thank Suzanne Fritz, Mary O’Connor, Bob O’Hara and Gergana Dasklova for comments on a previous version of the manuscript. DB, JSC, JH, SAB & MW were funded by the German Research Foundation (DB: Grant no BO 1221/23-1; JSC, JH, LMN, SAB & MW: via iDiv: FZT 118). SRS was funded by the National Science Foundation, USA (NSF 1400911). LHA was supported by Fundação para a Ciência e Tecnologia, Portugal (POPH/FSE SFRH/BD/90469/2012). ADB was supported by The Danish Council for Independent Research - Natural Sciences (DFF 4181-00565). MD was funded by the Scottish Funding Council (MASTS, grant reference HR09011) and MD & AEM by the ERC project BioTIME (250189) and BioCHANGE (727440). CW was supported by the Natural Environmental Research Council (grant number NE/L002531/).

## Author contributions

DB performed the analyses and wrote the first outline of the paper with AEB. All authors designed the study and helped draft the manuscript.

