## Supplementary material for "Mapping human pressures across the planet uncovers anthropogenic threat complexes": SOM

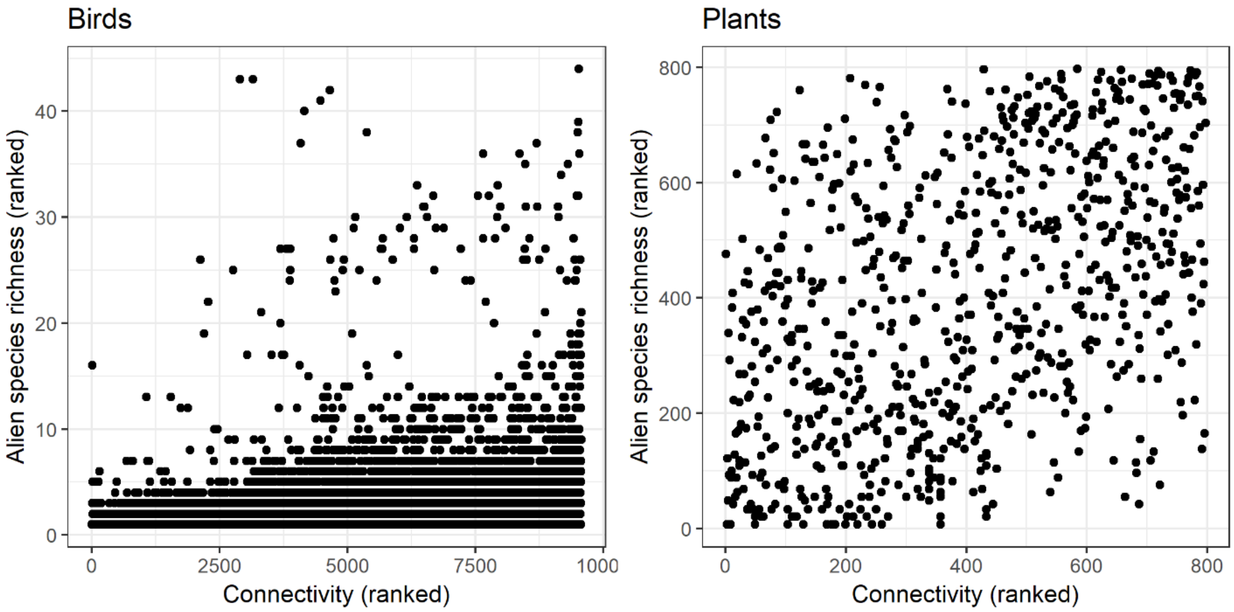

**Fig. S1** Spatial relationship between our proxy variable of invasive aliens (connectivity, based on transport infrastructure) and the number of alien species of birds and plants. For birds, a range map of alien species richness (Dyer *et al.* 2017) was intersected with our global grid of connectivity (resolution of 100 km). For plants, information on plant distributions were provided at regional scales (national, subnational or local scales) (van Kleunen *et al.* 2019), hence regional alien species richness was related to mean connectivity of each region.

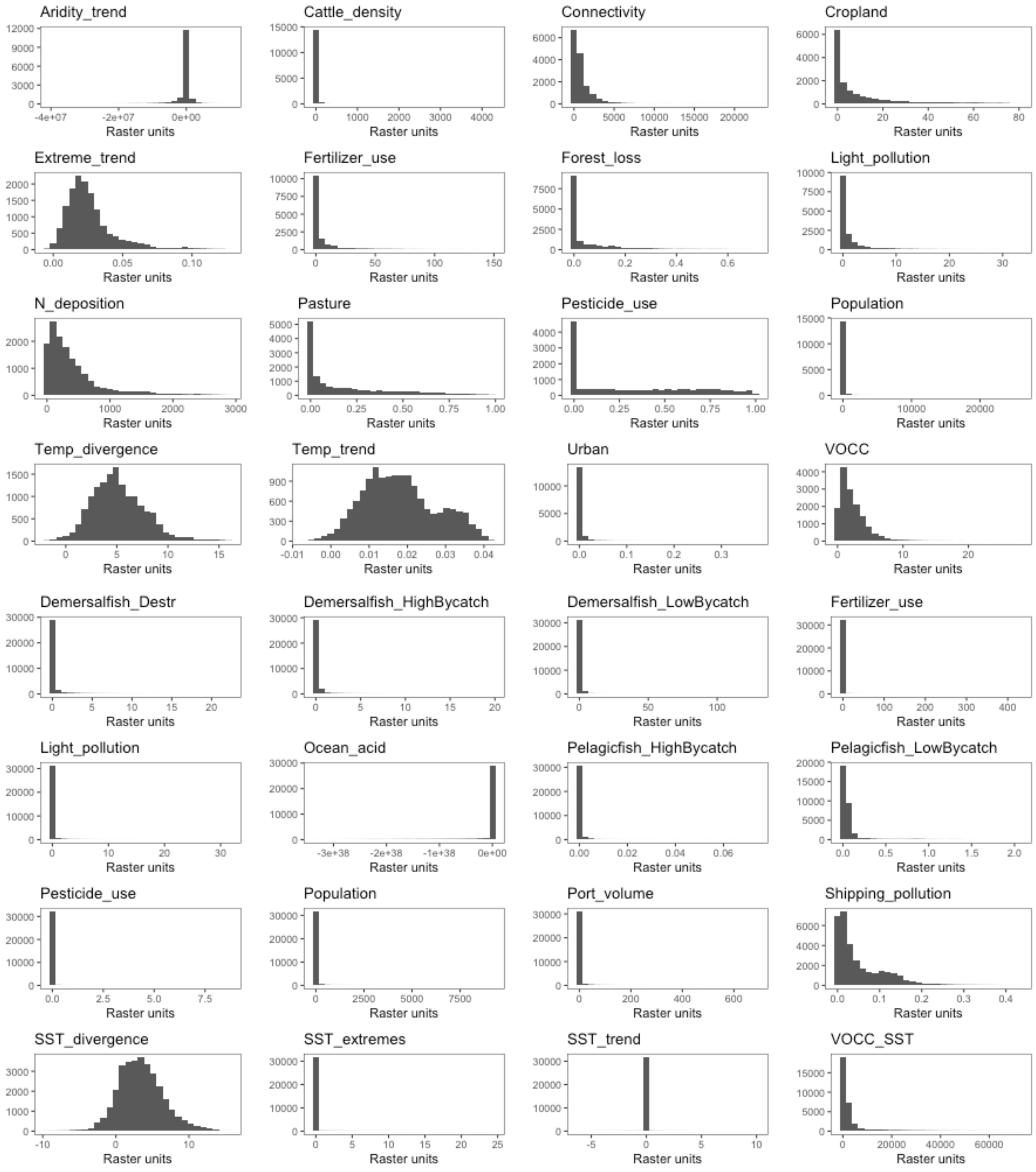

**Fig. S2** Distributions of the original values of each variable dataset. Because the datasets are highly skewed, we performed all analysis on the ranks of the values.

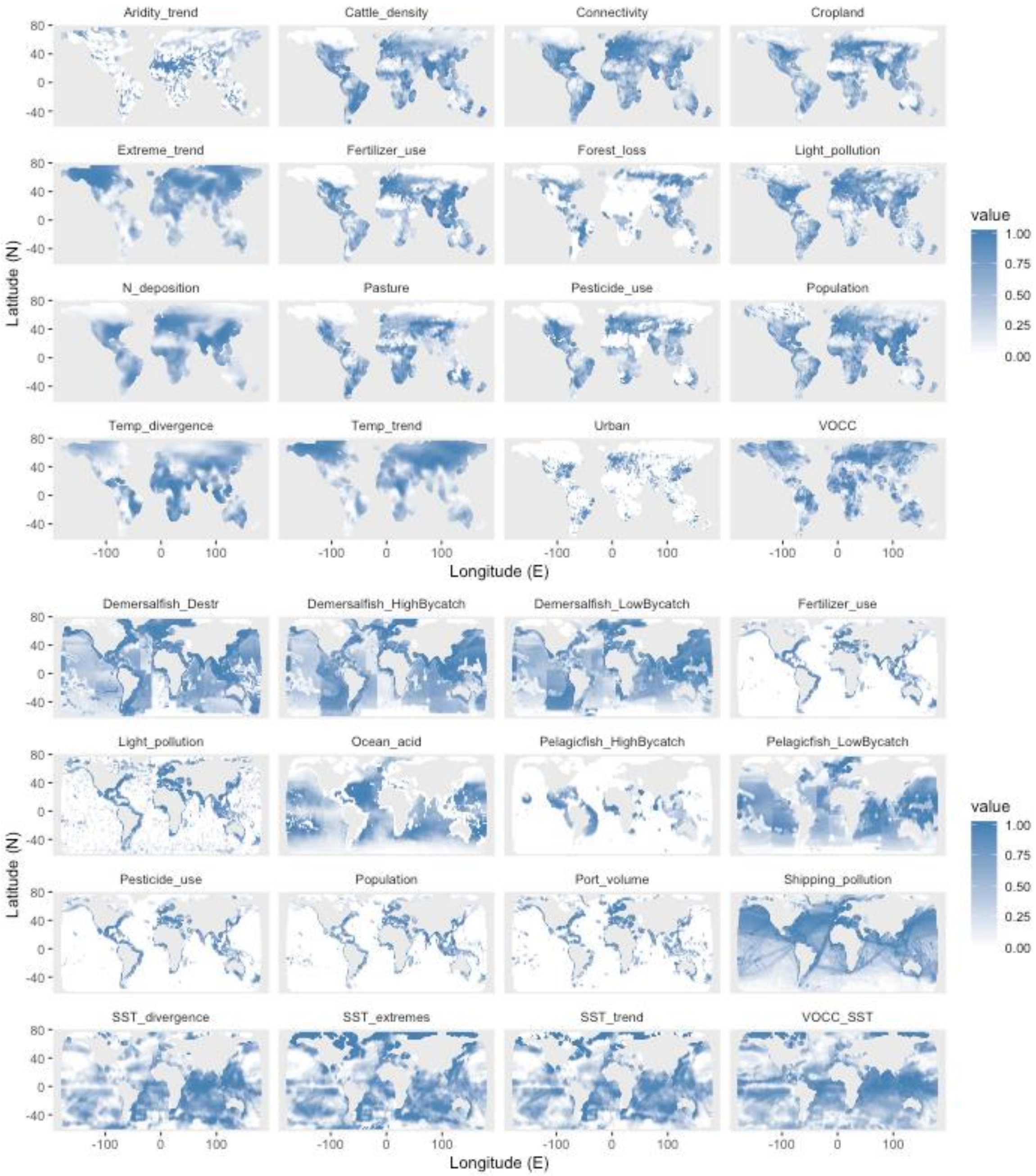

**Fig. S3** Global maps of each variable after ranking and scaling the values of each raster between 0 and 1.

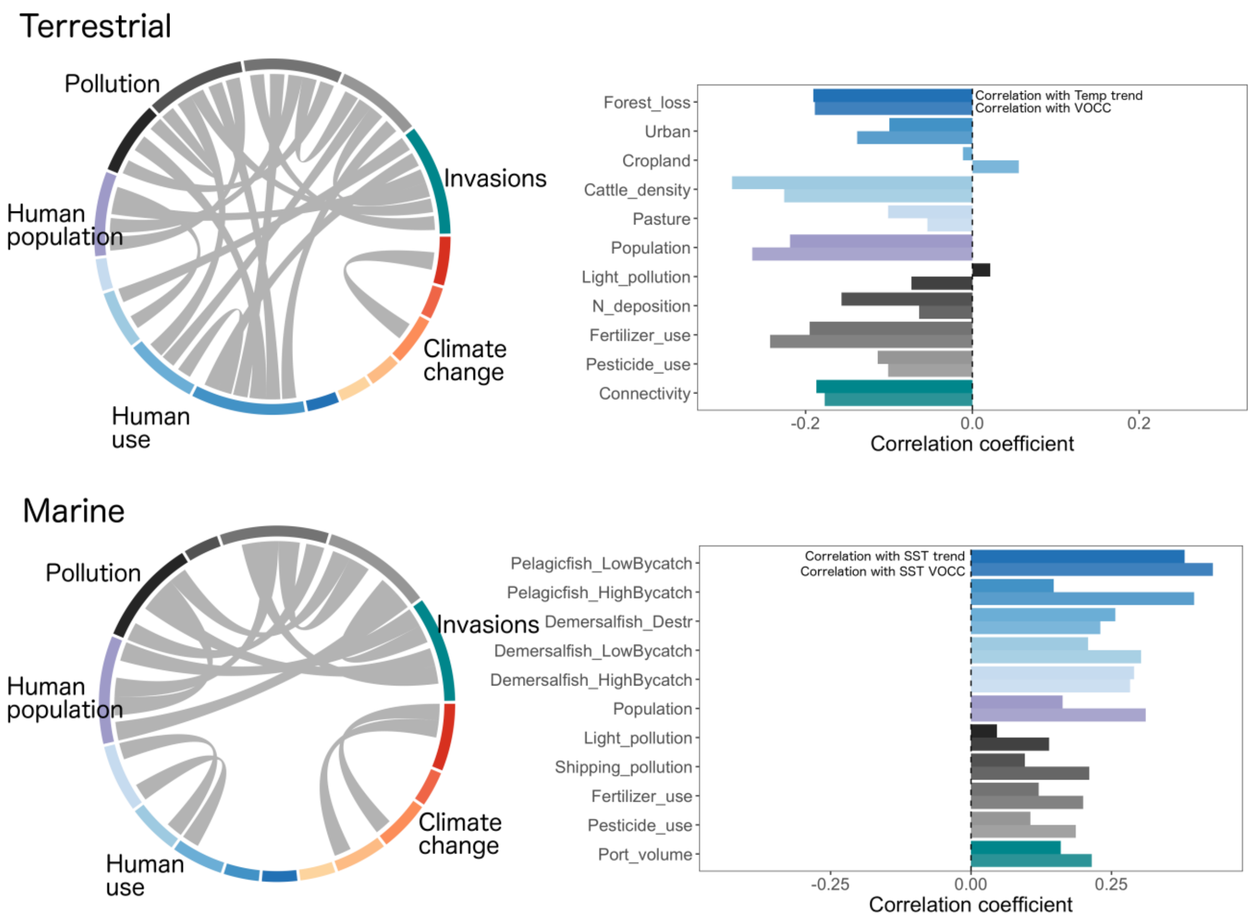

**Fig. S4** Analysis of correlations between variables on a grid of 500 km resolution. (Left) strong correlations (>0.7) among variables related to different drivers. (Right) correlations between each variable and temperature trend and velocity of climate change (VOCC). Colors refer to the same legend as Fig. 1 in the main text.

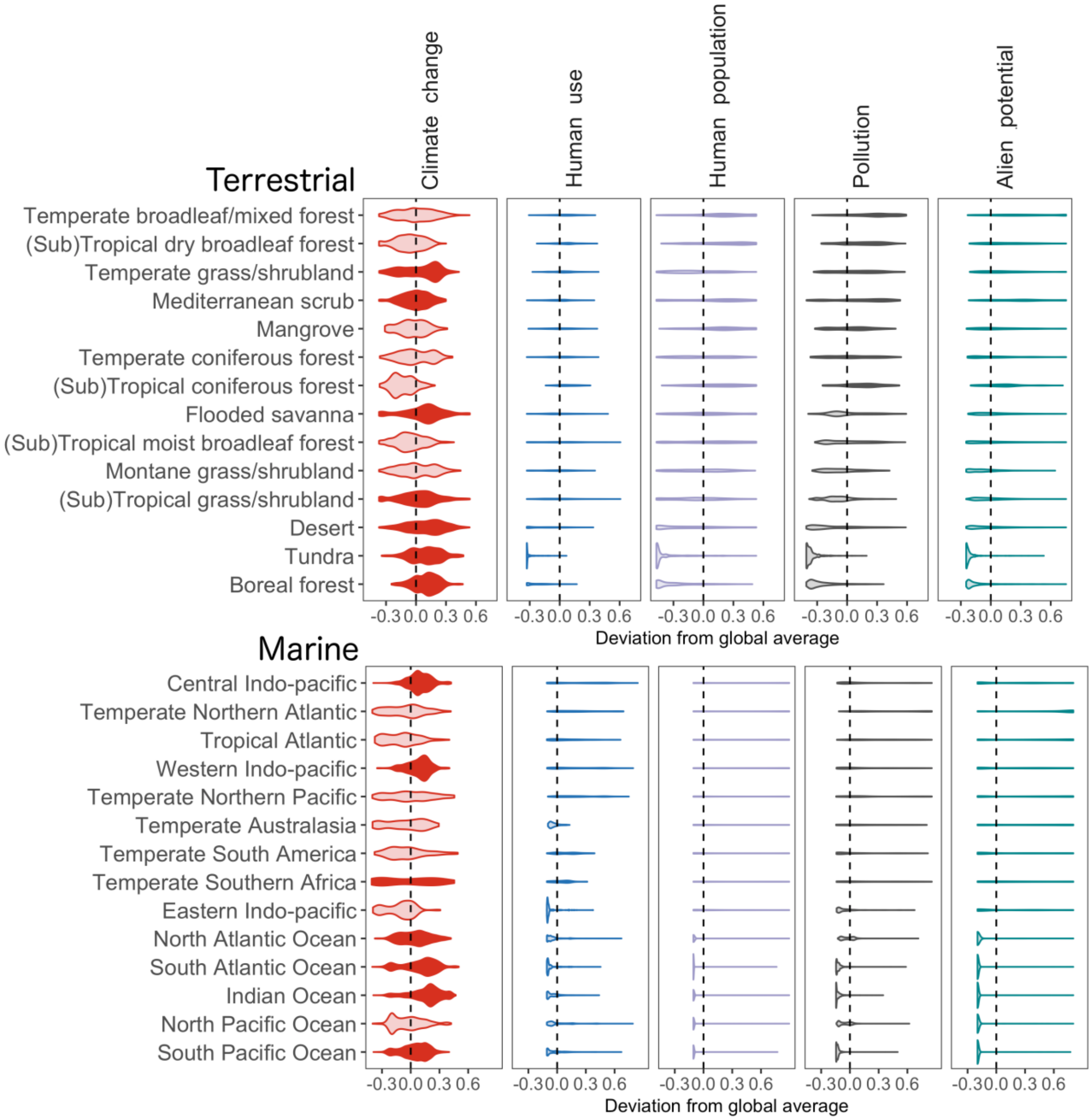

**Fig. S5**. Regions of the marine and terrestrial realms are exposed to distinct combinations of variables. As Fig. 3 except with different raster processing (logging rather than ranking dataset values). The violins show the distribution of values for each driver in each terrestrial and marine region. Violins with a median greater than the global median of each driver (centered on the dashed zero line) are colored in a darker color shade. Regions are presented in declining order of the sum of the driver means. Even when rasters are processed by logging the values, the distributions are still highly skewed; hence, we present in the main text the results when the rasters were processed by ranking the values.

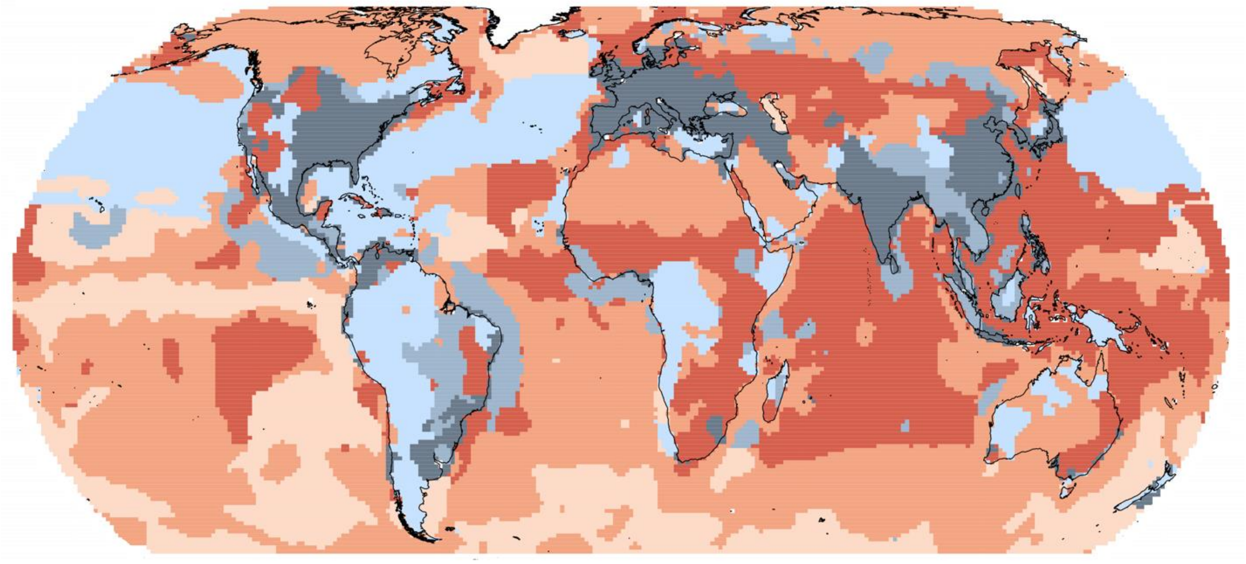

**Fig. S6** Different geographic regions of the world are exposed to different anthropogenic threat complexes. As Fig. 4 in the main text except with different raster processing (logging rather than ranking dataset values). These complexes were identified by their similarity of exposure to different variables on biodiversity. Each region is colored according to the same color scheme as Fig. 4 to facilitate comparison of the clustering.

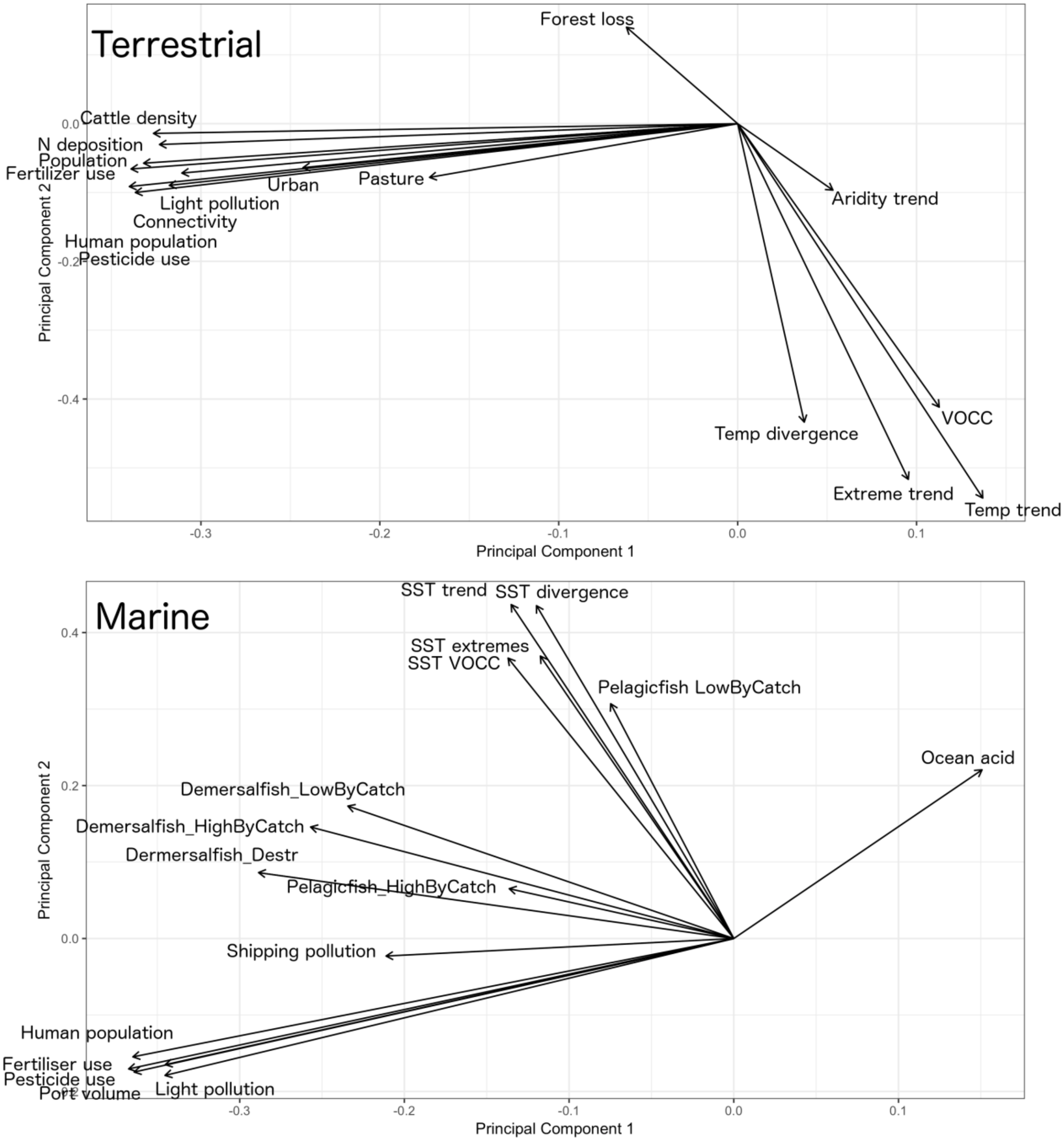

**Fig. S7** PCA biplots showing the correlations among variables in the terrestrial (top) and marine (bottom) realm. In both cases, the first two axes explained around 55% of the variation in the data

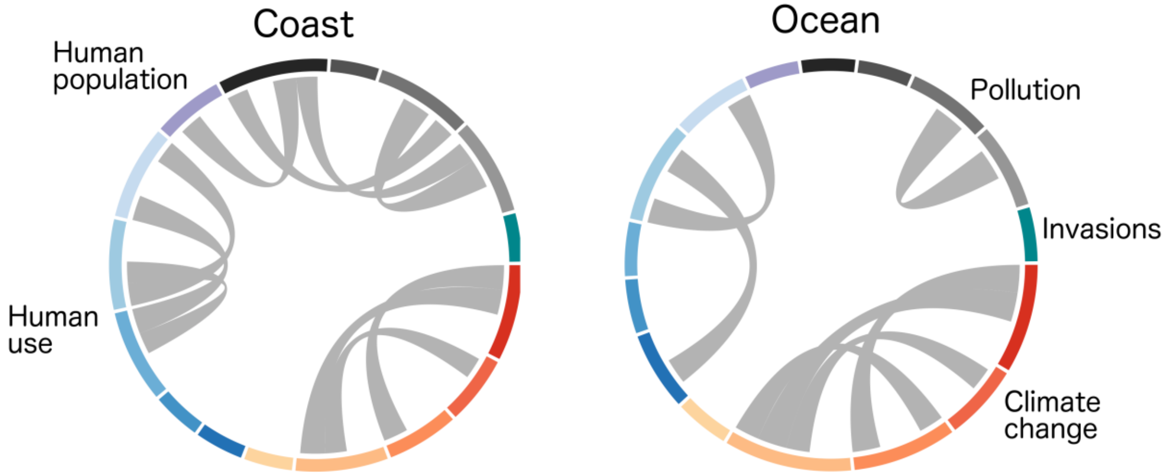

**Fig. S8** Correlations between variables for oceanic and coastal regions. We find fewer correlations between biodiversity change variables in oceanic regions than in coastal regions of the marine realm. Each link represents a significant relationship between two variables across 100 km grids covering each region. All links are positive correlations with strength >0.7 and were statistically significant after accounting for spatial autocorrelation. Colors refer to the same legend as in Fig. 1 in the main text.

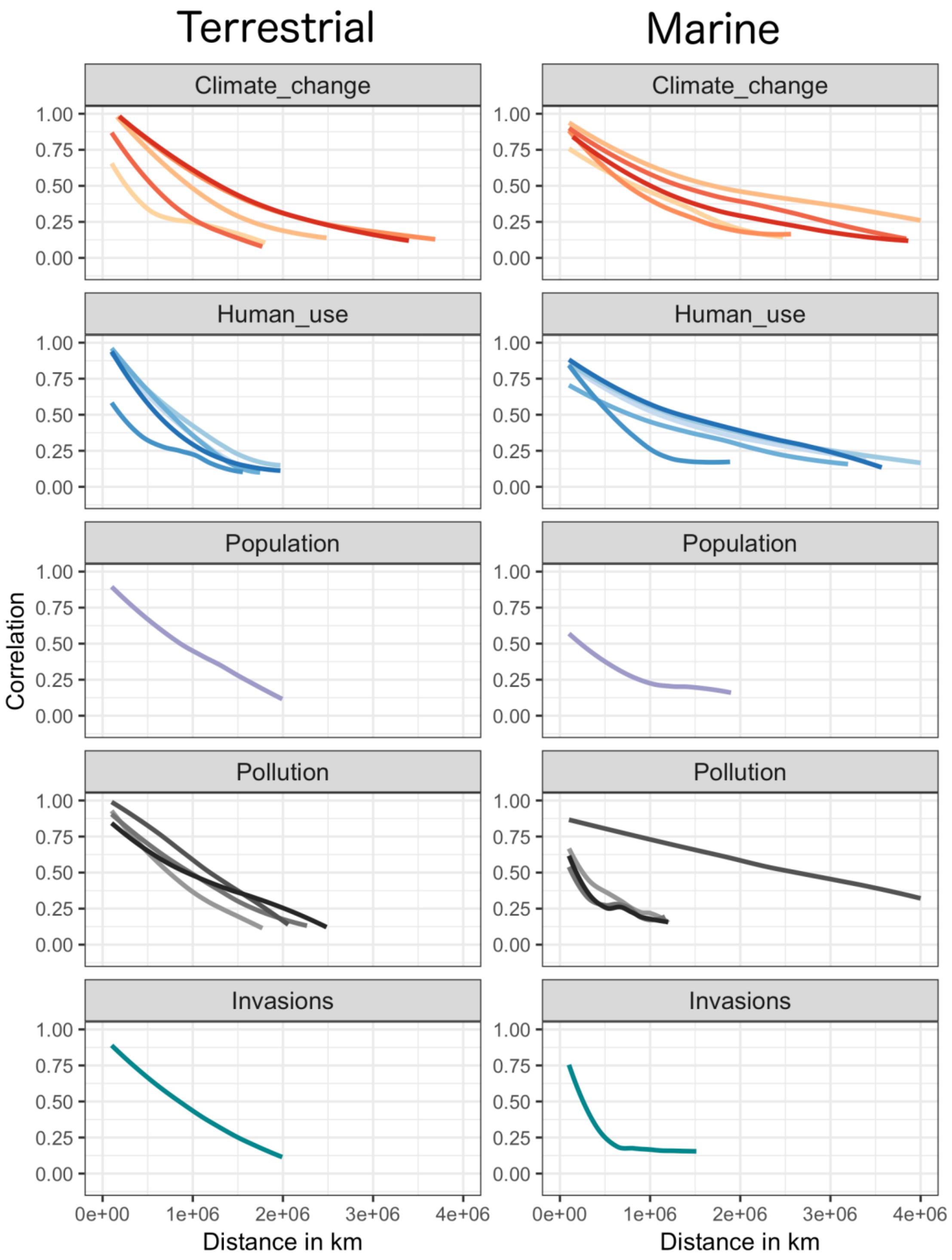

**Fig. S9** Correlograms showing the spatial autocorrelation (assessed by Moran's I) of the variables of each driver in each realm according to distance. Each line is plotted only as far as the extent of spatial autocorrelation remained statistically significant for five consecutive distance classes. Colors refer to the same legend as in Fig. 1 in the main text.

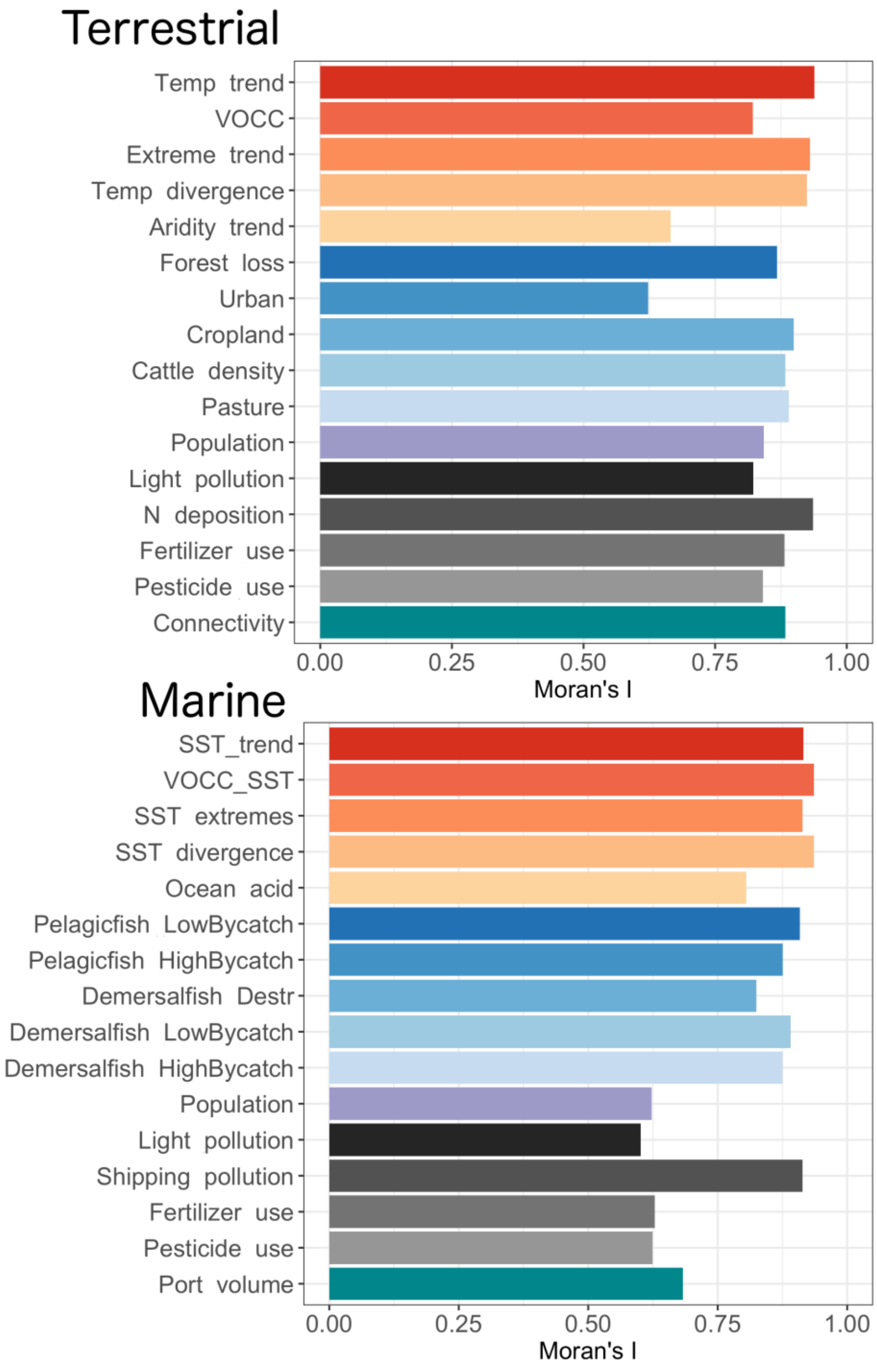

**Fig. S10** Estimates of Moran's I (larger values have greater spatial autocorrelation) for each variable in each realm.

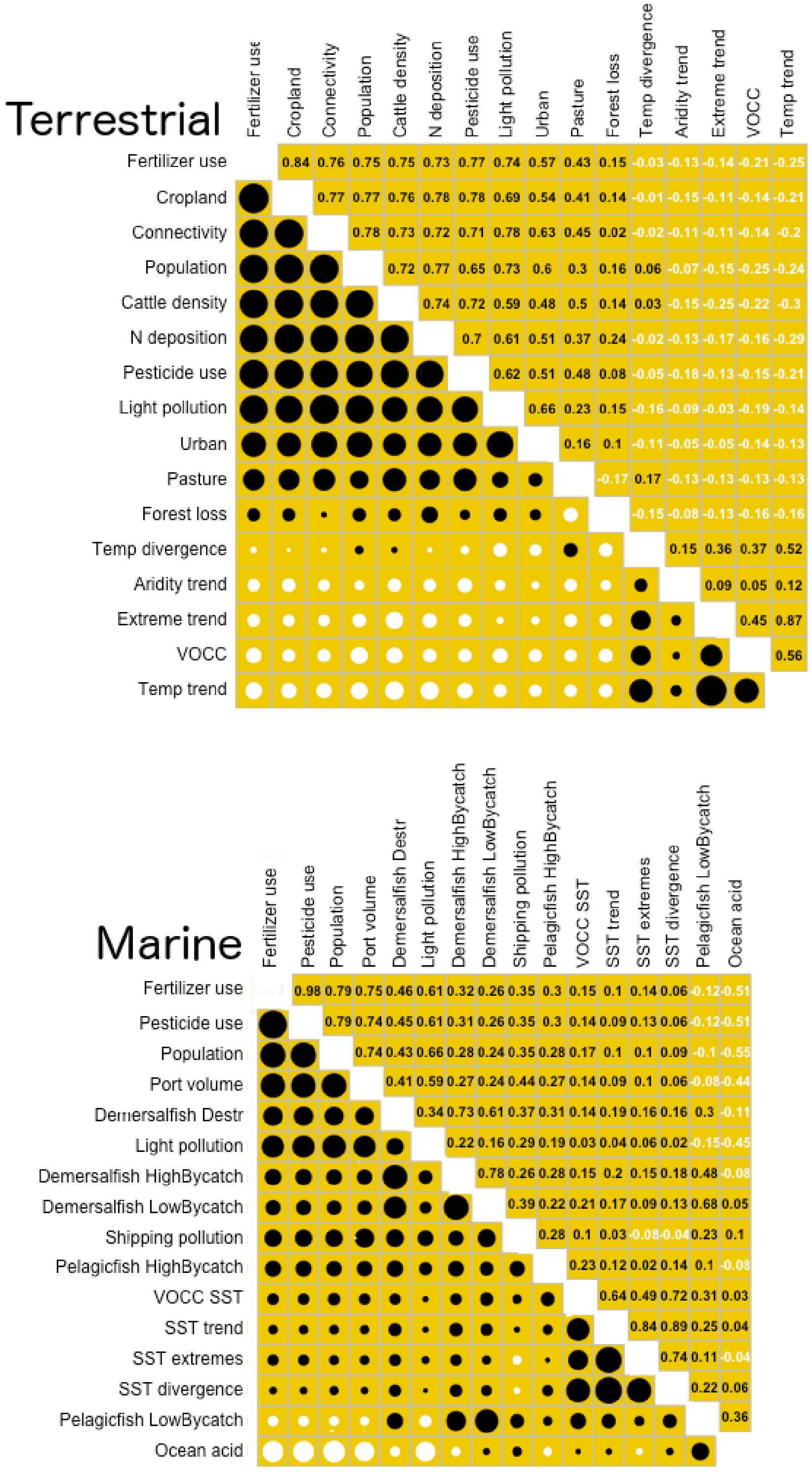

**Fig. S11** Correlations among variables at the global-scale in the terrestrial (top) and marine realms (bottom). Variables are ordered according to principal components (i.e., the most related variables are clustered together). Black circles indicate positive correlations and white circles negative ones; the size of the circles indicates the strength of the correlation.

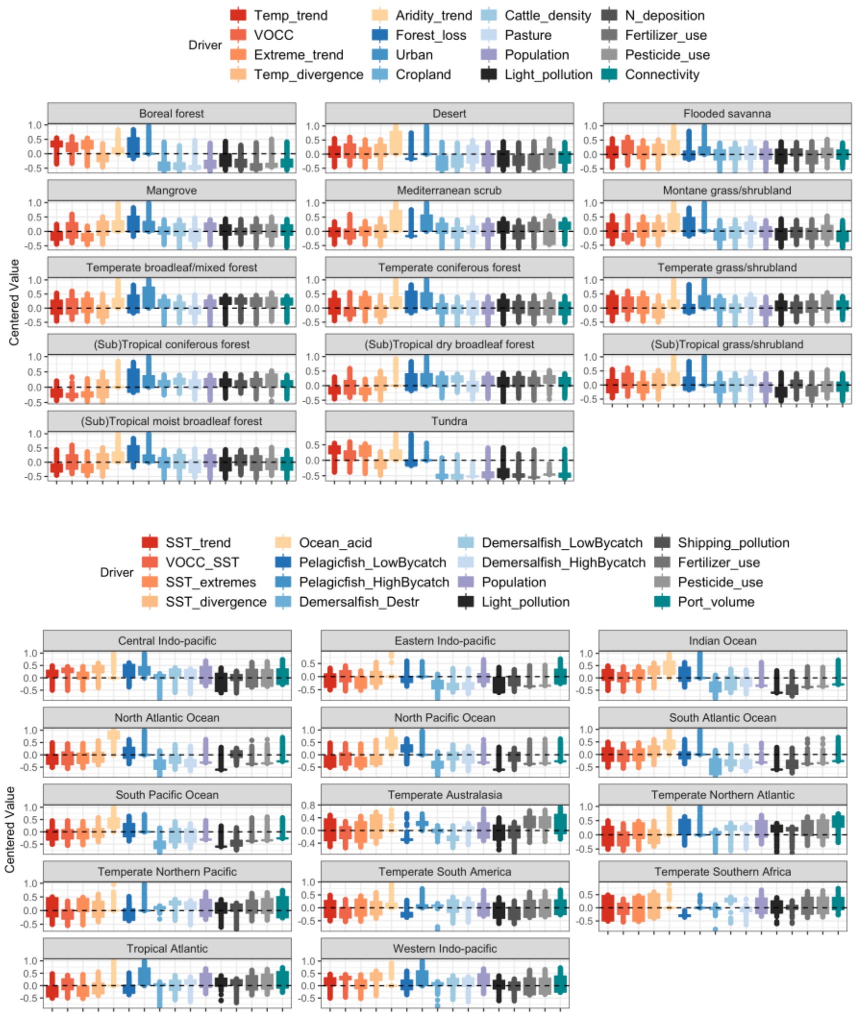

**Fig. S12** Distributions of the driver variables in each terrestrial and marine region. Each variable is centered by the average of that variable across all biomes, so that boxes above the dotted line represents greater than the average biome value and boxes below represent lower than the average. Boxes are delimited by the upper and lower quartiles; outliers beyond this are also shown.

The full names for the terrestrial regions are: Boreal forests / Taiga; Deserts and xeric shrublands; Flooded grasslands and savannas; Mangroves; Mediterranean Forests, woodlands and scrubs; Montane grasslands and shrublands; Temperate broadleaf and mixed forests; Temperate Coniferous Forest; Temperate grasslands, savannas and shrublands; Tropical and subtropical coniferous forests; Tropical and subtropical dry broadleaf forests; Tropical and subtropical grasslands, savannas and shrublands; Tropical and subtropical moist broadleaf forests; Tundra.

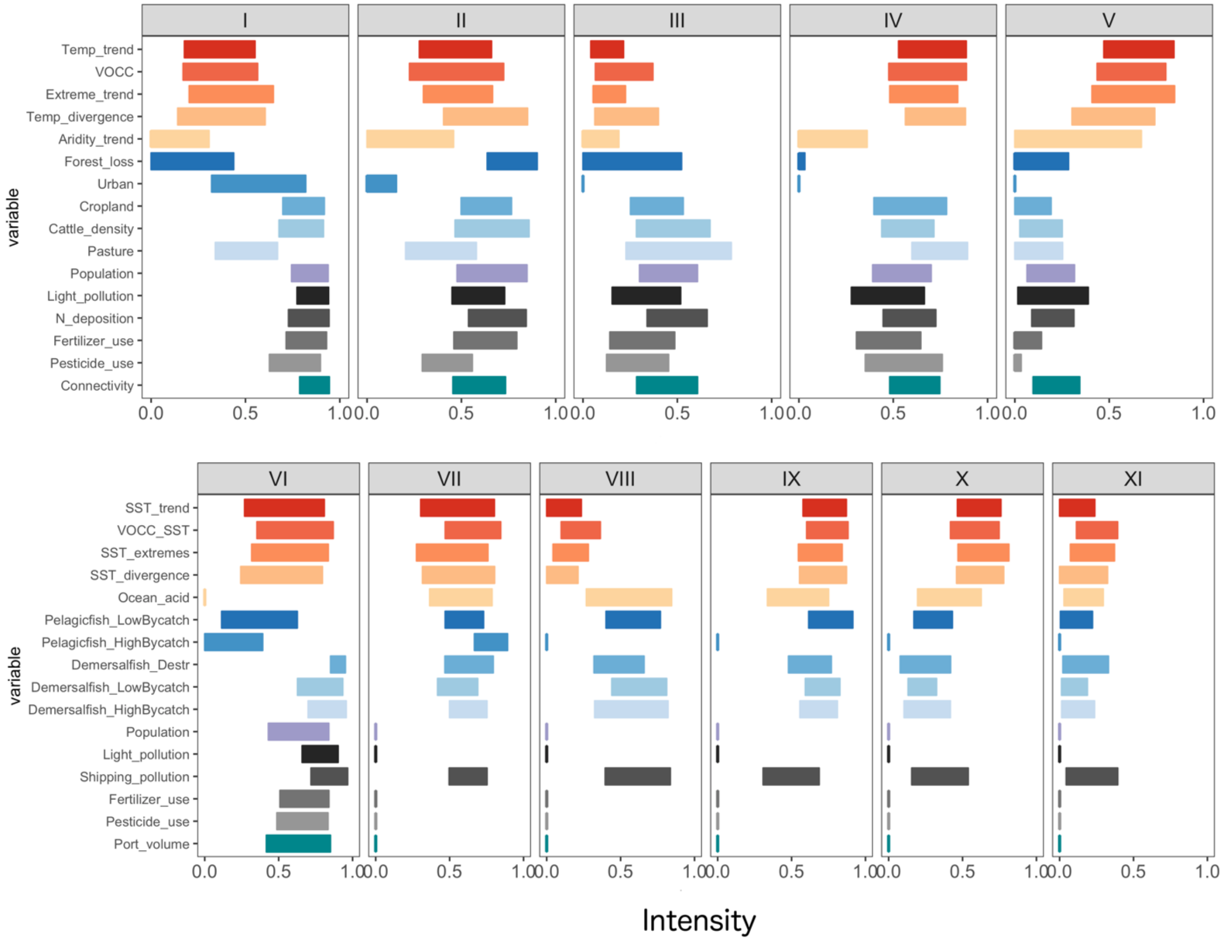

**Fig. S13** Anthropogenic threat complexes in the terrestrial (top row) and marine (bottom row) realms as shown in Fig. 4 in the main text. Each complex comprises differential exposure to a suite of variables. The crossbars showed the upper and lower quartiles of the intensity of each variable (scaled between 0 and 1).

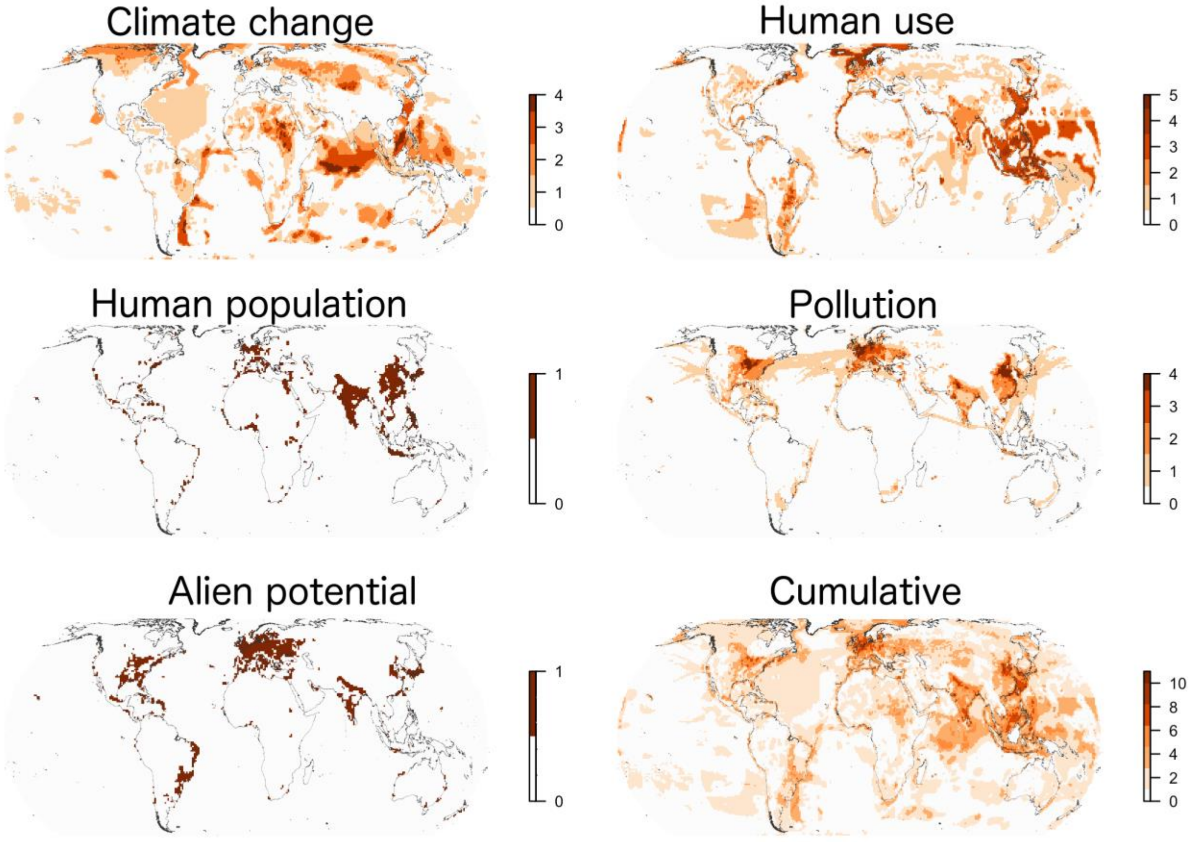

**Fig. S14** The number of the variables that each grid cell was in the highest 10% of values for each driver. Regions in the darkest orange are exposed to high intensities of multiple variables, while those in off-white are exposed to lower intensities of all (i.e., still potentially exposed to pressures but the magnitude of the pressures is not in the highest 10% of magnitude values). Note: Greenland was not included in the analysis due to missing data in several of the datasets.

**Table S1** Mapping of the driver variables included in our analysis in relation to the IUCN threat categories (Salafsky *et al.* 2008). We indicate where a spatial gridded data layer is "not available" or regarded as "not relevant" for a realm.

| IUCN threat categories | IUCN threat subcategories | Terrestrial (relevant layer) | Marine (relevant layer) | Comment |
|---|---|---|---|---|
| **Residential & commercial development** | Housing & urban areas | Urban | Not relevant | |
| | Commercial & industrial areas | Urban | Not relevant | |
| | Tourism & recreation areas | Urban | Not available | Not available in gridded form |
| **Agriculture & aquaculture** | Annual & perennial non-timber crops | Cropland | Not relevant | |
| | Wood & pulp plantations | Not available | Not relevant | not available globally: http://data.globalforestwatch.org/datasets/baae47df61ed4a73a6f54f00cb4207e0_5 |
| | Livestock farming & ranching | Pasture land; Cattle density | Not relevant | |
| | Marine & freshwater aquaculture | See comment | See comment | available for freshwater at riverthreat.net<br>not available in gridded form for marine |
| **Energy production & mining** | Oil & gas drilling | See comment | Not included | Available for the marine realm from Halpern et al. 2015 but not included since it was based on night lights, already included |
| | Mining & quarrying | Not available | Not available | Not available in gridded form |
| | Renewable energy | Not considered | Not considered | Not available in gridded form |

| | | | | |
|---|---|---|---|---|
| **Transportation & service corridors** | Roads & railroads | Connectivity | Not relevant | |
| | Utility & service lines | Not available | Not available | |
| | Shipping lanes | Not relevant | Correlated with shipping pollution | |
| | Flight paths | Not considered | Not considered | Not regarded as a large enough threat |
| **Biological resource use** | Hunting & collecting terrestrial animals | Not available | Not relevant | |
| | Gathering terrestrial plants | Not available | Not relevant | |
| | Logging & wood harvesting | Forest loss | Not relevant | |
| | Fishing & harvesting aquatic resources | See comment | Commercial fishing | available for freshwater at riverthreat.net |
| **Human intrusions & disturbance** | Recreational activities | Human population density | Human population density | |
| | War, civil unrest & military exercises | Not considered | Not considered | Regarded as too dynamic for inclusion |
| | Work & other activities | Not considered | Not considered | |
| **Natural system modifications** | Fire & fire suppression | Not considered | Not relevant | Not considered because difficult to link with human activities |
| | Dams & water management /use | See comment | Not relevant | available for freshwater at riverthreat.net |
| **Invasive & other problematic species, genes & diseases** | Invasive non-native/alien species/diseases | Connectivity | Port volume | |
| | Problematic native species/diseases | Connectivity | Port volume | |

| | | | | |
|---|---|---|---|---|
| | Introduced genetic material | Connectivity | Port volume | |
| **POLLUTION** | household sewage and urban waste water | Not available | Not available | |
| | Industrial & military effluents | Not available | Ships | |
| | agricultural and forestry effluents | Pesticide; Fertilizer | Pesticide; Fertilizer | |
| | garbage and solid waste | Not available | Not available | |
| | air-borne pollutants | Atmospheric nitrogen | Not considered | |
| | excess energy | Light pollution | Light pollution | |
| **GEOLOGICAL EVENTS** | Volcanoes | Not considered | Not considered | Not clearly linked to human activities |
| | Earthquakes /tsunamis | Not considered | Not considered | Not clearly linked to human activities |
| | Avalanches/l andslides | Not considered | Not considered | Not clearly linked to human activities |
| **CLIMATE CHANGE & SEVERE WEATHER** | Habitat shifting & alteration | Trend in temperature mean and divergence | Trend in temperature mean and divergence | |
| | Droughts | Aridity trend | Not relevant | |
| | Temperature extremes | Trends in extreme temps | Trends in extreme temps | |
| | Storms & flooding | Not considered | Not considered | |
| | Other | | Ocean acidification | |

**Table S1** Metadata and web links for each variable dataset used in the analysis

| Variable | Realm | Best Data Layer | Time series | Resolution | Description/Url/Reference |
|---|---|---|---|---|---|
| **Temperature** | T | CRU v 4.02 | Yes | 0.5° | mean monthly and yearly temperatures (°C) https://crudata.uea.ac.uk/cru/data/hrg/ (Harris *et al.* 2014) |
| **Aridity change** | T | CRU v 4.02 | Yes | 0.5° | ratio of mean monthly and yearly pet (mm $day^{-1}$) and precipitation (mm) https://crudata.uea.ac.uk/cru/data/hrg/ (Harris *et al.* 2014) |
| **Sea surface temperature** | M | HadISST | Yes | 1° | mean monthly and yearly sea surface temperatures (°C) https://www.metoffice.gov.uk/hadobs/hadisst/data/download.html (Rayner *et al.* 2003) |
| **Ocean acidification** | M | Ocean Acidification | Yes* (2000-2009 vs 1870) | 1 $km^2$ | change in aragonite saturation state https://www.nceas.ucsb.edu/globalmarine/impactbyactivity (Halpern *et al.* 2008) |
| **Pasture** | T | Pasture fraction | No (2000) | 5' | fraction of cell area (0-1) based on agricultural inventory data and satellite-derived land cover data http://www.earthstat.org/ (Ramankutty *et al.* 2008) |
| **Cropland** | T | Cropland fraction | No (2005) | 5' | fraction of cell area (0-1) based on national and subnational agricultural data and satellite-derived land cover data (Fritz *et al.* 2015) |
| **Cattle density** | | Gridded Livestock of the World | No (2005) | 1 km | FAOSTAT national estimates and modelled downscaling (Robinson *et al.* 2014) |
| **Forest loss** | T | Land-Use Harmonization 2 (primary forest cover) | Yes | 0.25° | fraction of cell area (0-1) using FAO national wood harvest volume data and an ecosystem model |

| | | | | | |
|---|---|---|---|---|---|
| | | | | | http://luh.umd.edu/<br>(Hurtt *et al.* (in prep)) |
| **Urban cover** | T | MODIS | No (2001) | 5' | Urban cover (0 or 1) based on satellite-derived land cover data<br>http://glcf.umd.edu/data/lc/<br>(Friedl *et al.* 2010) |
| **Fishing** | M | Commercial fishing layers | No (1999-2003) | 1 $km^2$ | tons of caught fish per ton of carbon<br>https://www.nceas.ucsb.edu/globalmarine/impactbyactivity<br>(Halpern *et al.* 2008) |
| **Population density** | T | SEDAC population data v4 | No (2000) | 30” | UN-adjusted population density<br>http://sedac.ciesin.columbia.edu/data/set/gpw-v4-population-density/data-download<br>(Center for International Earth Science Information Network - CIESIN - Columbia University 2017) |
| **Coastal population** | M | Coastal population | No (1992-2002) | 1 $km^2$ | number of people within 25 km radius<br>https://www.nceas.ucsb.edu/globalmarine/impactbyactivity<br>(Halpern *et al.* 2008) |
| **N deposition** | T | Atmospheric nitrogen deposition | No (1993) | 5º x 3.75º | mg $N/m^2$ of total inorganic nitrogen (N), NHx (NH3 and NH4+), and NOy<br>http://webmap.ornl.gov/ogcdown/dataset.jsp?ds_id=830<br>(Dentener 2006) |
| **Fertilizer application** | T | Nitrogen fertilizer application (v1) | No (1994-2001) | 0.5° | kg of Nitrogen fertilizer per hectare of cropland<br>http://sedac.ciesin.columbia.edu/data/set/ferman-v1-nitrogen-fertilizer-application<br>(Potter *et al.* 2010) |
| **Pesticides** | T | Riverthreat.net: Pesticide loading | No (2000) | 0.5° | kg of pesticide per hectare of cropland<br>http://www.riverthreat.net/data.html<br>(Vorosmarty *et al.* 2010) |

| | | | | | |
|---|---|---|---|---|---|
| **Light pollution** | T/M | NOACC NGDC stable night lights | No (2006) | 1 km | radiance values https://knb.ecoinformatics.org/#view/doi:10.5063/F15718ZN (Halpern *et al.* 2008) |
| **Coastal pollution** | M | Pesticide, Fertilizer | No (1993-2002) | 1 $km^2$ | average annual use in agricultural land https://knb.ecoinformatics.org/#view/doi:10.5063/F15718ZN (Halpern *et al.* 2008) |
| **Shipping pollution** | M | Shipping pollution | No (2004-2005) | 1 $km^2$ | ship activity (number of ships) https://knb.ecoinformatics.org/#view/doi:10.5063/F15718ZN (Halpern *et al.* 2008) |
| **Invasions** | T | Accessibility (Travel time) | No (2000) | 30” | travel time to major cities (in hours and days) http://forobs.jrc.ec.europa.eu/products/gam/ (Nelson 2008) |
| **Invasions** | M | Port volume (cargo volume at ports) | No (1999-2003) | 1 $km^2$ | amount of cargo traffic at ports https://knb.ecoinformatics.org/#view/doi:10.5063/F15718ZN (Halpern *et al.* 2008) |

*These data had already been calculated as a change between the two time points by the data provider.
